# Nanobody-guided proximity proteomics reveals conformationally encoded molecular neighborhoods of LRRK2

**DOI:** 10.64898/2026.11.23.766131

**Authors:** Benjamin Riebenbauer, Felix Hoffmann, Christian Galicia, Alessandro Ragazzini, Francesco Raimondi, Karsten Boldt, Wim Versées, Christian Johannes Gloeckner

**Author notes:** Corresponding author: Christian Johannes Gloeckner.

## Abstract

Leucine-rich repeat kinase 2 (LRRK2) is a multidomain signaling protein implicated in Parkinson’s disease whose activity is closely linked to conformational state. How these structural transitions shape the cellular molecular environment of LRRK2 remains poorly understood. Here, we combined nanobody-mediated conformational control with miniTurbo proximity proteomics to define molecular neighborhoods associated with inhibitory, neutral, and activating LRRK2 states. Structural modeling suggested that the activating nanobody Nb22 targets the WD40 domain and destabilizes interactions maintaining the autoinhibited conformation. Consistent with this model, activation was accompanied by ordered, reciprocal remodeling of the LRRK2 proximity proteome, with progressive enrichment of centrosomal, ciliary, and RAB/endolysosomal machinery and depletion of soluble cytosolic, proteostasis-associated and 14-3-3 proteins. Notably, the Nb22-associated proteome combined centrosomal and endolysosomal components previously linked to distinct LRRK2 signaling environments. These findings support a model in which conformational transitions reorganize the spatial molecular neighborhood of LRRK2 and thereby couple kinase state to distinct cellular signaling networks.

## Introduction

Dominant pathogenic variants in the *LRRK2* gene represent the most common genetic cause of familial Parkinson’s disease (PD), and genetic variation at this locus also contributes to the risk of sporadic PD (Paisan-Ruiz *et al*, 2004; Simon-Sanchez *et al*, 2009; Zimprich *et al*, 2004). *LRRK2* encodes a large multidomain signaling protein containing tandem ROC-COR GTPase and serine/threonine kinase domains. A major function of LRRK2 is the phosphorylation of a subset of Rab G-proteins that regulate membrane trafficking pathways (Alessi & Pfeffer, 2024). Consequently, LRRK2 has emerged as a major therapeutic target and several kinase inhibitors are currently under clinical development (Kluss et al., 2022). Beyond its established role in PD, however, LRRK2 is increasingly recognized as a multifunctional signaling protein involved in the regulation of vesicular trafficking, lysosomal homeostasis, cytoskeletal dynamics and ciliogenesis, and inflammatory responses (Alessi and Sammler, 2018). Structurally, LRRK2 comprises a complex arrangement of protein–protein interaction and catalytic domains, including N-terminal armadillo, ankyrin, and leucine-rich repeat domains, a ROC-COR tandem G-protein module, a serine/threonine kinase domain, and a C-terminal WD40 domain (Guaitoli *et al*, 2016). A growing body of evidence indicates that LRRK2 function is closely linked to its spatial organization within the cell. Particularly, activated LRRK2 accumulates at specific membrane compartments and phosphorylates a subset of Rab G-proteins that regulate endolysosomal trafficking (Bonet-Ponce *et al*, 2020; Lee *et al*, 2020; Rinaldi *et al*, 2023; Steger *et al*, 2016). Furthermore, LRRK2 has been implicated in cytoskeletal dynamics and ciliogenesis, while altered LRRK2 activity has also been associated with defects in centrosomal cohesion (Meixner *et al*, 2011; Ordóñez *et al*, 2019; Tombesi *et al*, 2026). Recent cryo-EM studies have revealed that LRRK2 adopts multiple conformational states characterized by dynamic interdomain interactions and varying degrees of structural compaction (Deniston *et al*, 2020; Myasnikov *et al*, 2021; Villagran Suarez *et al*, 2026; Zhu *et al*, 2023). These structural transitions are increasingly recognized as important determinants of LRRK2 activity, localization and cellular function (Schmidt *et al*, 2021). Emerging evidence suggests that distinct conformational states may preferentially engage specific cellular compartments and protein interaction networks. Collectively, these findings suggest that LRRK2 functions within specialized molecular neighborhoods whose composition is likely dictated by both subcellular localization and conformational state. However, how distinct structural states are translated into specific interaction networks remains poorly understood.

Despite the recent elucidation of an active structure upon RAB29 binding, current experimental approaches provide only limited opportunities to dissect the relationship between LRRK2 conformation and cellular organization (Zhu *et al*., 2023). Indeed, while recent structural studies have begun to define inactive and active-like LRRK2 conformations (Villagran Suarez *et al*., 2026; Zhu *et al*., 2023), experimental tools capable of selectively stabilizing these states in living cells remain limited. In our recent proximity proteomic study, we have demonstrated that LRRK2 activity states are associated with distinct molecular environments, including centrosomal, ciliary, vesicular, and endolysosomal protein networks (Eckert *et al*, 2026). However, although LRRK2-selective type I inhibitors stabilize an active-like kinase-domain conformation, how discrete conformational and catalytic states shape the surrounding protein network remains unresolved.

Nanobodies provide a unique opportunity to address this challenge. Due to their small size and high binding specificity, nanobodies can selectively stabilize defined conformational states of target proteins. We previously identified a panel of LRRK2-directed nanobodies that exert distinct functional effects, including inhibitory, neutral, and activating activities (Singh *et al*, 2022). These reagents enable direct manipulation of the LRRK2 conformational landscape independently of conventional pharmacological approaches and therefore provide powerful tools to investigate how structural state influences the organization of LRRK2-associated signaling networks.

We hypothesized that structural transitions within LRRK2 function as molecular switches that selectively recruit specific protein assemblies, thereby generating spatially distinct signaling environments. To test this model, we combined nanobody-mediated conformational control with miniTurbo-based proximity proteomics to define interaction networks associated with inhibitory, neutral, and activating LRRK2 states. Comparative analysis revealed extensive rewiring of the LRRK2 proximity proteome, including ordered remodeling of functional protein modules across nanobody-defined states. Activating LRRK2 states preferentially engaged centrosomal, ciliary, vesicular and endolysosomal networks, accompanied by reciprocal changes in proteostasis-associated molecular environments.

## Results

### Selecting suitable affinity probes that stabilize defined LRRK2 activity states

To investigate how distinct LRRK2 conformations shape the molecular environment of the kinase, we established a proximity-labeling workflow combining miniTurbo-mediated biotinylation with conformation-selective affinity probes (Figure 1A). We previously generated and characterized a panel of LRRK2-specific nanobodies that bind LRRK2 with nanomolar affinity and differentially modulate its kinase activity through allosteric mechanisms (Singh *et al*., 2022). Based on their functional properties, we hypothesized that these nanobodies stabilize distinct conformational states of LRRK2 and therefore provide molecular tools to interrogate conformation-dependent changes in the LRRK2 proximity proteome.

**Figure 1:**
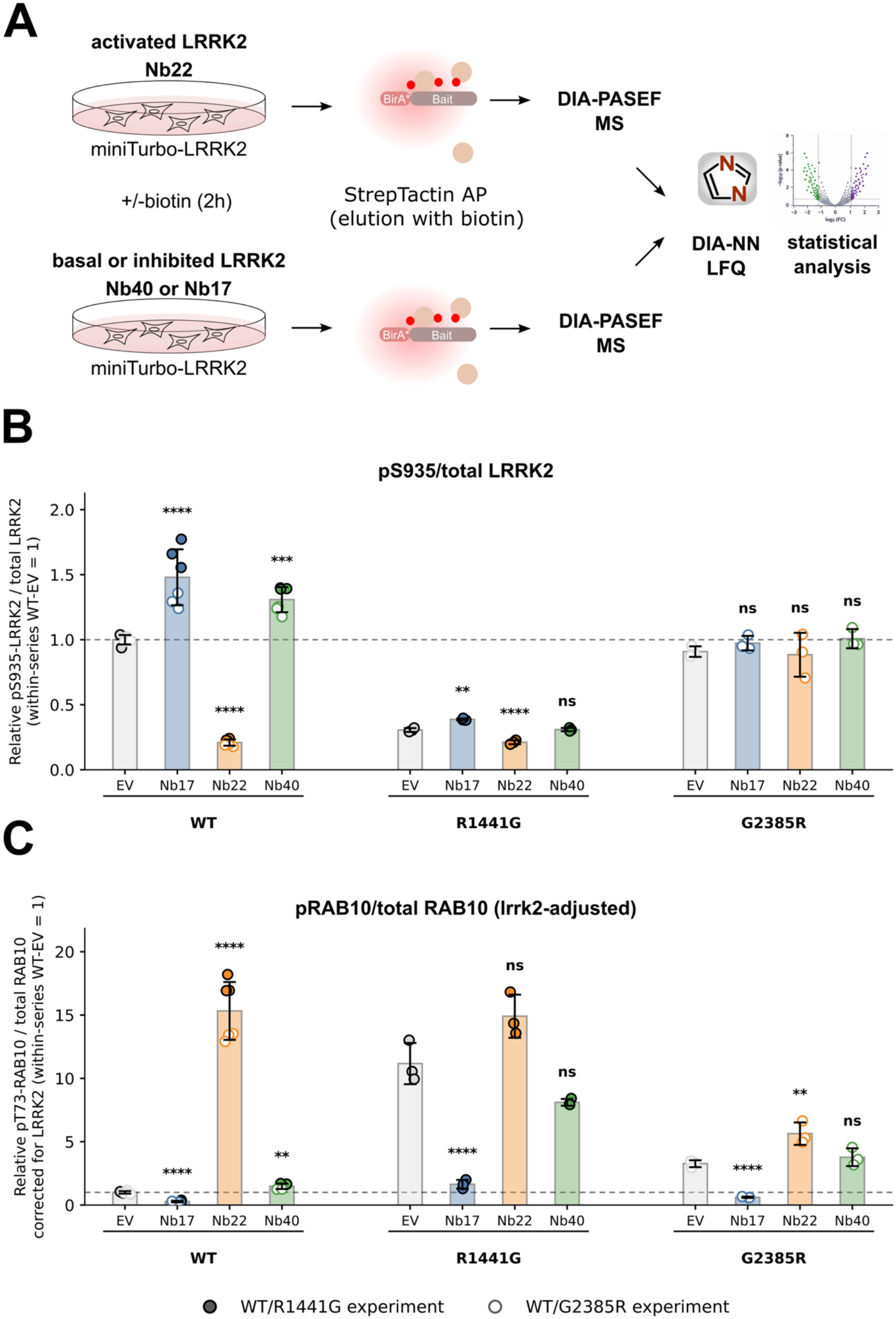
Nanobody-dependent modulation of LRRK2 activity and conformational biomarkers. (A) Experimental workflow for miniTurbo proximity labeling using miniTurbo-LRRK2 together with the activating nanobody Nb22 or the inhibitory/neutral nanobodies Nb17 and Nb40. (B) Quantitative capillary immunoblot analysis of pS935-LRRK2 normalized to total LRRK2. Values were normalized to the WT-EV control within each experimental series (WT = 1). Individual biological replicates are shown as circles, with filled and open symbols indicating the independent WT/R1441G and WT/G2385R experimental series, respectively. Horizontal lines indicate the mean and error bars represent SD. (C) Quantification of pT73-RAB10 normalized to total RAB10 and corrected for total LRRK2 abundance. Values were normalized to the WT-EV (empty vector) control within each experimental series (WT = 1). Individual biological replicates are shown as circles, with horizontal lines indicating the mean and error bars representing SD. Statistical significance in both panels (B, C) was determined by genotype-wise Dunnett-adjusted multiple comparisons versus the EV control following within-series normalization. ns, P ≥ 0.05; P < 0.05 (*); P < 0.01 (**); P < 0.001 (***); P < 0.0001 (****). Each mutant condition comprises three biological replicates. WT data from the WT/R1441G and WT/G2385R experimental series were combined after normalization to the corresponding within-series WT-EV control (WT = 1), yielding six observations for each WT condition.

From this study, we have selected three nanobodies, including a neutral, an inhibitory and an activating binder: Nb40, Nb17 and Nb22, respectively. Nb22 and Nb40 both recognize the C-terminal WD40 domain but differ markedly in their effects on LRRK2 activity, allowing conformational changes to be investigated while minimizing epitope-dependent effects. The inhibitory Nb17 was selected because it presumably engages multiple LRRK2 domains, as suggested by earlier CL-MS (chemical crosslinking combined with mass spectrometry) experiments and by the observation that efficient binding was only detected for full-length LRRK2 among the constructs tested.

We next validated these nanobodies in cell-based activity assays (Figure 1B and C). Total LRRK2 expression varied considerably between conditions. We therefore additionally normalized the pT73-RAB10/total RAB10 ratio to β-actin-corrected total LRRK2 abundance. Nb17 robustly inhibited RAB10 phosphorylation by more than 70% when co-expressed with either WT or R1441G LRRK2. Because the residual pRAB10 signal approached the limit of quantification, the sequential normalization to total RAB10 and LRRK2 likely inflated the apparent residual signal. Accordingly, the remaining kinase activity is likely overestimated in this case (Supplementary Figure 1A). For transparency, and because normalization of pT73-RAB10 to total RAB10 represents the standard analysis in the field, a side-by-side comparison to the singly normalized data is provided in Supplementary Figure 1B. The corresponding total LRRK2 expression levels normalized to β-actin are shown in Supplementary Figure 1C. In WT LRRK2, Nb22 increased RAB10 phosphorylation approximately fifteenfold (approximately fourfold when no correction for LRRK2 abundance is used). In cells expressing the pathogenic R1441G variant, basal RAB10 phosphorylation was increased approximately tenfold relative to wild-type LRRK2, consistent with the hyperactive nature of this variant. Nb17 strongly suppressed R1441G LRRK2 activity, whereas Nb22 did not further increase pRAB10 levels, suggesting that R1441G LRRK2 may already operate close to the maximal activity detectable in this cellular assay. Nb40 did not produce a consistent directional effect across genotypes, supporting its classification as a largely neutral binder. It modestly increased LRRK2-normalized RAB10 phosphorylation in the WT background but reduced it in the R1441G background.

In addition to RAB phosphorylation, we determined the phosphorylation state of S935, a well-established marker site for N-terminal LRRK2 phosphorylation and 14-3-3 binding (Dzamko *et al*, 2010; Gloeckner *et al*, 2010). This site is rapidly dephosphorylated upon treatment with type I LRRK2 inhibitors, but not type II inhibitors, and has therefore been proposed as a conformational/regulatory state biomarker (Alessi & Pfeffer, 2024; Dzamko *et al*., 2010; Raig *et al*, 2025). Consistent with this concept, Nb22 induced a marked reduction in S935 phosphorylation in WT LRRK2. Conversely, Nb17 increased S935 phosphorylation in WT LRRK2, consistent with Nb17 favoring a more autoinhibited conformational state. In contrast, R1441G showed strongly reduced S935 phosphorylation across conditions, and Nb17 did not restore S935 phosphorylation to WT levels despite strongly suppressing RAB10 phosphorylation. These findings indicate that catalytic output toward RAB substrates and phosphorylation of the N-terminal biomarker site S935 can become uncoupled, particularly in the context of the hyperactive R1441G variant. These results are also consistent with a recent preprint providing an extended cell-based analysis of the LRRK2 nanobody set we previously developed (Ordóñez *et al*, 2026).

### Structural models suggest distinct mechanisms of Nb22, Nb40, and Nb17 action

To investigate the structural basis for the distinct effects of Nb22, Nb40, and Nb17 on LRRK2 kinase activity, we generated AlphaFold3 models of the corresponding nanobody–LRRK2 complexes (Figure 2A; Supplementary Figure 2). Although AlphaFold3 substantially improves the prediction of antibody-antigen interactions compared with previous versions, we limited the modeling to the smallest LRRK2 constructs that had been experimentally validated to bind the respective nanobodies efficiently in our previous study (Singh et al., 2022). Model assessment was based on AlphaFold3 interface confidence metrics (ipTM/PAE), agreement with previously reported DSSO crosslinking constraints (Singh *et al*., 2022), and structural compatibility with the cryo-EM structure of autoinhibited LRRK2 (PDB: 7LHW) (Myasnikov *et al*., 2021). While these computational models need further experimental validation, and the overall pose confidence was higher for Nb40 than for Nb22, both models were sufficiently supported to enable hypothesis generation. The final models for the WD40 binders Nb22 and Nb40 are shown in Figure 2A.

**Figure 2:**
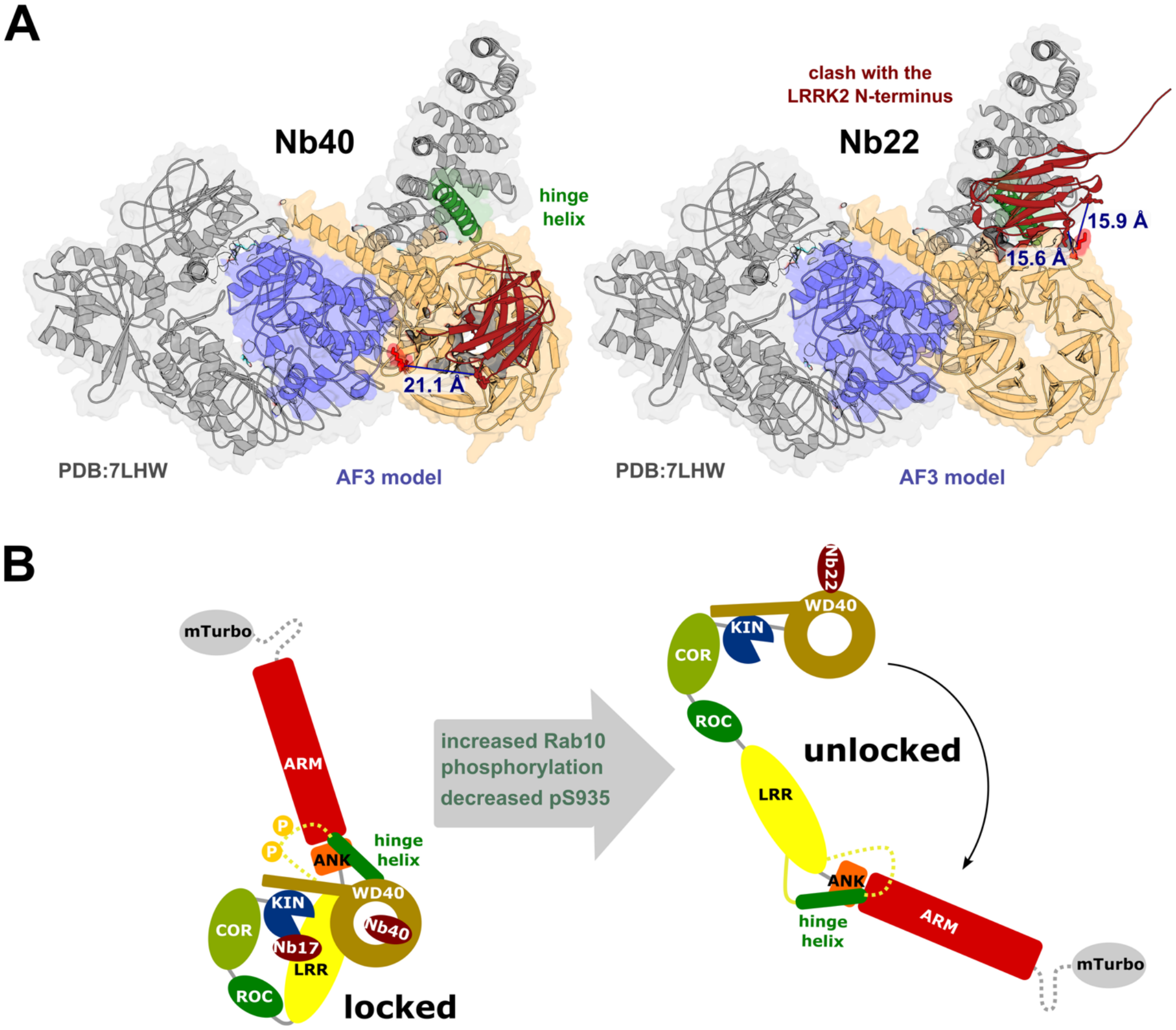
LRRK2 conformational states controlled by nanobodies. (A) AlphaFold3 models for the complex of Nb40 (left) and Nb22 (right) with the Kinase-WD40 (the minimum binding epitope previously identified for these nanobodies (Singh *et al*., 2022)). The models were superimposed onto the full-length autoinhibited LRRK2 structure 7LHW. While the pose of Nb40 is in good agreement with the autoinhibited state, the pose of Nb22 causes clashes with the N-terminal Armadillo and the LRR domain, particularly the hinge helix, consistent with stabilization of an unlocked conformation. The poses of both nanobodies are in good agreement with DSSO crosslinks previously determined by XL-MS (Singh *et al*., 2022). (B) Mechanistic models for the inactive (locked) vs active (unlocked) LRRK2 conformation. This model is in good agreement with recent EM structures (Villagran Suarez *et al*., 2026). The predicted binding sites of selected nanobodies are indicated.

Interestingly, the AlphaFold3 models suggest that Nb22 binds to an epitope within the WD40 domain located in close proximity to the previously identified hinge helix, which resides in an extended loop of the N-terminal portion of the LRR domain. Superposition of the Nb22-bound model onto the full-length autoinhibited LRRK2 structure (PDB: 7LHW) reveals steric clashes between Nb22 and the hinge helix, suggesting that Nb22 binding may displace the N-terminal LRR domain from its wrapped conformation around the kinase domain. Such a rearrangement would be consistent with disruption of the autoinhibited state. Both the top-ranking Nb22 and Nb40 models are in good agreement with our previous DSSO crosslinking data (Singh *et al*., 2022). In the Nb22 model, Nb22 K45 and K67 form crosslinks with LRRK2 K2382, with Cα–Cα distances of 15.9 Å and 15.6 Å, respectively. In addition, this model agrees well with recent structural analyses of autoinhibited and active conformations of monomeric LRRK2 reported by Villagran Suarez et al. (2026). Notably, the Nb22 docking pose overlaps with the position of the G2385R PD risk variant, which has been reported to increase RAB10 phosphorylation in cells (Kalogeropulou *et al*, 2022; Nguyen *et al*, 2026; Tezuka *et al*, 2022). We therefore asked whether Nb22 is impaired in its ability to further increase the kinase activity of the moderately hyperactive G2385R variant. In cell-based assays, G2385R caused an approximately threefold increase in RAB10 phosphorylation compared with wild-type LRRK2. Whereas Nb22, tested in parallel, increased RAB10 phosphorylation by approximately one order of magnitude above wild-type baseline, it produced only a modest increase in the presence of G2385R and failed to elevate RAB10 phosphorylation to the levels observed for Nb22-bound wild-type LRRK2. These findings are consistent with the proposed binding model, in which the G2385R substitution impairs productive Nb22 engagement. Notably, in agreement with the RAB10 phosphorylation data and in contrast to the pronounced dephosphorylation of pS935 observed upon Nb22 binding to wild-type LRRK2, Nb22 did not induce a detectable reduction in pS935 phosphorylation in the G2385R background (Figure 1B and C, third panel).

In contrast, the AlphaFold3 model of the Nb40–WD40 complex is fully compatible with the autoinhibited LRRK2 structure, providing a structural rationale for the largely neutral behavior of this nanobody. According to the model, Nb40 binds within the central pore of the WD40 β-propeller without perturbing the intramolecular interactions that stabilize the autoinhibited conformation. In our previous study, we identified one crosslink between Nb40 and LRRK2, which is fully satisfied by the model, with a C-alpha distance of 21.1 Å between LRRK2 K2485 and Nb40 K77. Thus, both Nb22 and Nb40 models are in excellent agreement with the distance constraints previously obtained by XL-MS (Singh *et al*., 2022).

Compared with Nb22 and Nb40, the binding mode of Nb17 was predicted with lower confidence by AlphaFold3. In the selected AF3-derived model, Nb17 binds predominantly to the C-terminal LRR/LRR–ROC linker surface, with additional contacts to the C-terminal lobe of the kinase domain. The pose is compatible with the previously reported CL-MS data: the crosslinks between Nb17 K64 and LRRK2 K1290, K1316 and K2078 correspond to Cα– Cα distances of 18.3, 29.2 and 34.3 Å, respectively, all within the relaxed 35 Å threshold commonly applied to DSSO crosslinks when conformational dynamics are considered (Supplementary Figure 2). K1290 and K1316 are located in the LRR–ROC linker, a structured element in autoinhibited LRRK2 that has been proposed to become disordered during activation (Deyaert *et al*, 2019; Galicia *et al*, 2024; Weng *et al*, 2023). The resulting composite LRR–kinase interface is also consistent with the observed requirement for full-length LRRK2 for high-affinity Nb17 binding. The model therefore provides a structural hypothesis in which Nb17 restricts rearrangement of the LRR– ROC linker and kinase-proximal region, thereby favoring a closed, autoinhibited LRRK2 conformation and limiting access of bulky Rab substrates to the kinase domain.

In conclusion, the biochemical and structural data support a model in which Nb22 activates LRRK2 by destabilizing its autoinhibited conformation. Specifically, Nb22 binding appears to displace the N-terminal LRR region from its inhibitory interaction with the kinase domain, thereby facilitating adoption of an active conformation. A mechanistic scheme of the two states is shown in Figure 2B.

### Nanobody-stabilized conformational state-dependent interactomes of LRRK2

To investigate how nanobody-defined LRRK2 states influence the local molecular environment of the kinase, we co-expressed the inhibitory nanobody Nb17, the activating nanobody Nb22, or the neutral nanobody Nb40 together with miniTurbo (mT)-LRRK2 and performed proximity labeling in the presence and absence of biotin (Figure 1A). Quantitative proximity proteomics identified robust sets of biotinylated proteins associated with each nanobody condition. Overall, the miniTurbo datasets largely recapitulated the interaction networks identified in our previous proximity proteomic analyses while additionally revealing a number of candidate interactors, likely reflecting the increased sensitivity and proteome coverage afforded by data-independent acquisition (DIA) mass spectrometry. In particular, we have identified a series of RAB and RAB-effector proteins, including well-established LRRK2 effectors (Supplementary Figure 3A-C).

To delineate state-specific molecular environments of LRRK2, we compared proteins that were significantly enriched in biotin-capture across the three nanobody conditions. This analysis revealed 100, 87, and 107 proteins that were exclusively enriched with Nb17, Nb40, and Nb22, respectively (Supplementary Fig. 3D). Over-representation analysis of the functionally opposing Nb17 and Nb22 conditions produced largely distinct annotation profiles (Supplementary Fig. 3E). Proteins specific to Nb17 were enriched for nucleoplasmic localization, protein-modification processes, Parkinson’s disease, and neurodevelopmental delay. Conversely, Nb22-specific proteins were associated with cytoplasmic and cytoskeletal compartments, including the Rho GTPase cycle, the MAD1L1–MAD2L1 complex, the ciliary landscape, and dynein-binding intraflagellar-transport proteins. These data indicate that nanobody-mediated modulation of LRRK2 establishes distinct molecular environments, particularly involving ciliary organization and intracellular transport in the activating Nb22 condition.

To determine whether these distinct functional signatures were also reflected in quantitative differences in the LRRK2 proximity proteome, we next directly compared protein abundances across the three nanobody conditions, restricting the analysis to biotin-treated samples. This approach minimized potential distortions arising from non-biotinylated background proteins and enabled a more direct assessment of nanobody-dependent remodeling of the LRRK2 molecular environment. Comparative analysis revealed pronounced remodeling of the LRRK2 molecular neighborhood upon expression of the activating nanobody Nb22. Both the Nb22 versus Nb40 (Figure 3A-C, Supplementary Figure 4) and Nb22 versus Nb17 (Figure 4A-C, Supplementary Figure 5) comparisons identified highly overlapping sets of significantly enriched proteins, indicating that Nb22 induces a proximity proteome that is clearly distinct from both neutral and inhibitory LRRK2 states.

**Figure 3:**
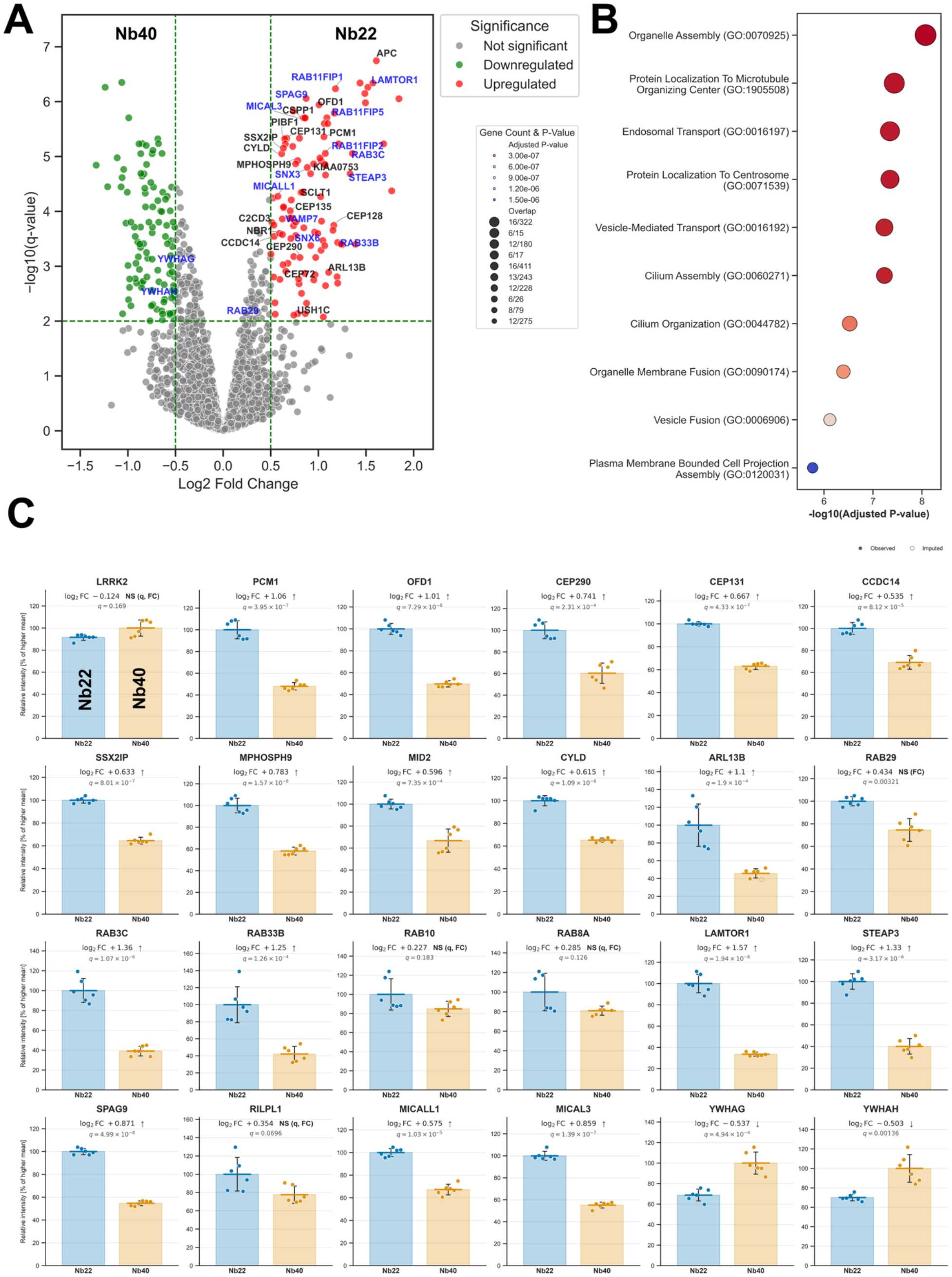
Rewiring of the LRRK2 proximity proteome by the activating nanobody Nb22 compared to the largely neutral nanobody Nb40. (A) Volcano plot showing differential protein abundance analyzed using a limma-like moderated t-test with multiple imputation and Rubin pooling; P values were adjusted using the Benjamini– Hochberg method (N=6 [2 biological and 3 technical replicates]; Rubin MI; FDR=0.01, N proteins=4035). (B) GO-term analysis of the proximity proteome enriched with Nb22. (C) Pairwise comparison of selected proteins (mean normalized abundance N=6 [2 biological and 3 technical replicates]), error bar= SD, enriched protein set to 100%, LIMMA q-value is shown (ns= non-significant, up and down arrows indicate the direction of significant differences).

**Figure 4:**
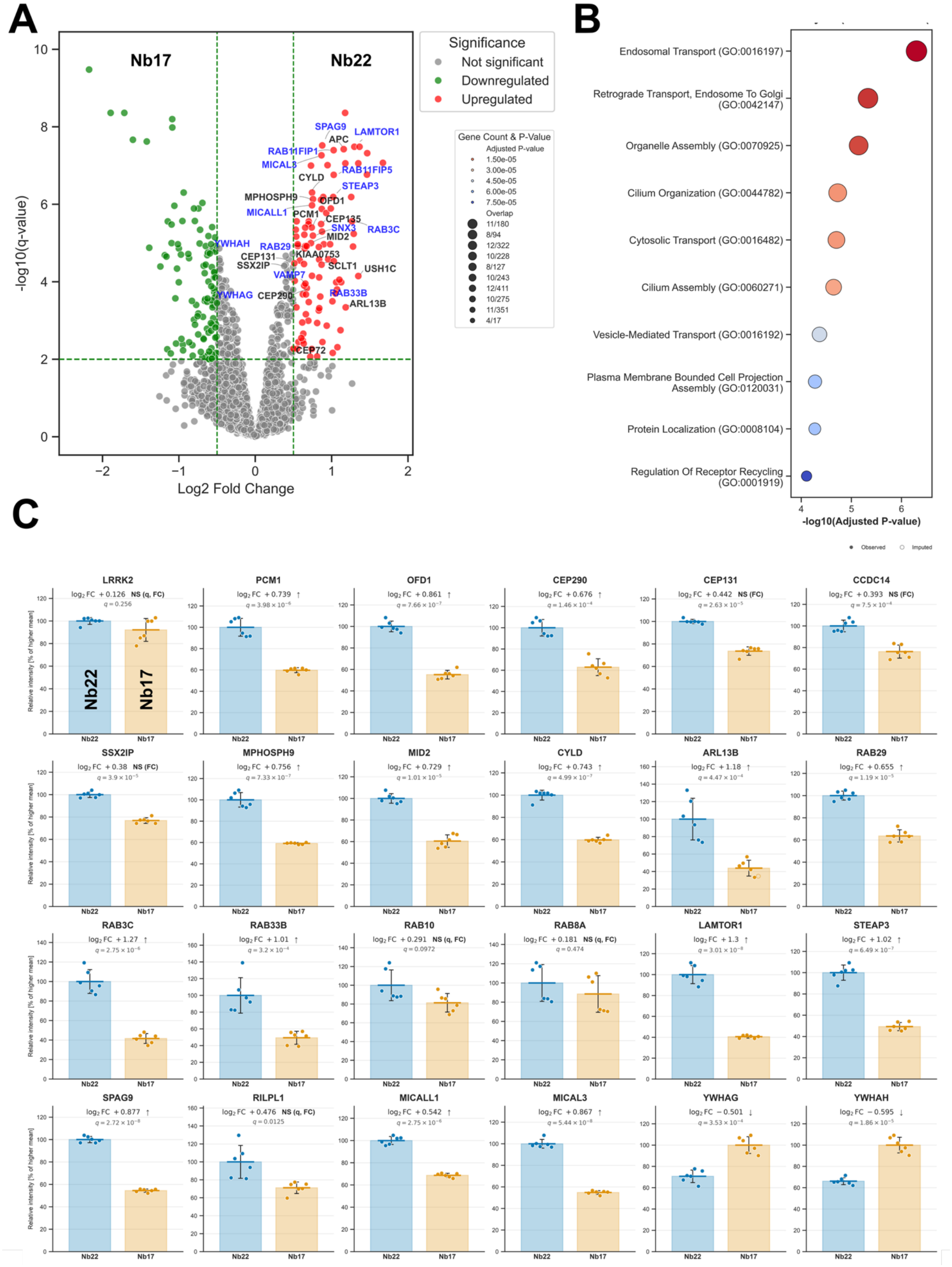
Rewiring of the LRRK2 proximity proteome by the activating nanobody Nb22 compared to the inhibitory nanobody Nb17. (A) Volcano plot showing differential protein abundance analyzed using a limma-like moderated t-test with multiple imputation and Rubin pooling; P values were adjusted using the Benjamini–Hochberg method (N=6 [2 biological and 3 technical replicates]; Rubin MI; FDR=0.01, N proteins=4011). (B) GO-term analysis of the proximity proteome enriched with Nb22. (C) Pairwise comparison of selected proteins (mean normalized abundance N=6 [2 biological and 3 technical replicates]), error bar= SD, enriched protein set to 100%, LIMMA q-value is shown (ns= non-significant, up and down arrows indicate the direction of significant differences).

Functional annotation of the significantly enriched proteins revealed prominent signatures associated with centrosomal and ciliary biology, vesicle trafficking, membrane organization, and cytoskeletal processes (Figure 3B, Figure 4B). Notably, proteins enriched in the Nb22 datasets overlapped with proteins previously identified in two distinct LRRK2 signaling environments: the proximity proteome induced by type I kinase inhibition and the interactome associated with RAB29-mediated activation of LRRK2 (Eckert *et al*., 2026).

To enable a direct comparison between nanobody-induced and inhibitor-induced proximity proteomes, we repeated the inhibitor experiments using the same magnetic bead-based enrichment workflow and DIA acquisition strategy employed for the nanobody datasets. This approach minimized technical differences between datasets and facilitated a robust assessment of shared and unique features of the respective LRRK2 signaling environments (Supplementary Figure 6-9). Consistent with our previous analyses, significant remodeling of the proximity proteome was primarily observed following treatment with the type I inhibitor MLi-2. The MLi-2-induced interactome was characterized by enrichment of numerous proteins associated with the basal body, primary cilium, and centriolar satellites, confirming the strong centrosomal and ciliary signature associated with inhibitor-stabilized LRRK2.

Comparison of the MLi-2 and Nb22 datasets revealed extensive overlap between the proteins enriched under both conditions. These shared proteins included numerous centrosomal and ciliary components, suggesting recruitment of related molecular assemblies. Among the most prominently enriched proteins were several centriolar satellite proteins, including the core scaffold PCM1 as well as OFD1, SSX2IP and CEP131 (Figure 3C, 4C, Supplementary Figure 6C, 8C). Additional shared proteins included components of the basal body and primary cilium, further supporting the strong centrosomal and ciliary signature observed under both conditions.

Notably, the deubiquitinase CYLD, a regulator of centriolar satellite proteostasis (Douanne *et al*, 2019) and which we recently identified as a candidate LRRK2 effector (Eckert *et al*., 2026), is significantly enriched in the Nb22-associated proximity proteome. The convergence of conformation-selective proximity proteomic and co-evolutionary analyses on CYLD further supports a functional association between CYLD and activated LRRK2 signaling states. Consistent with this notion, CYLD co-expression significantly increased the steady-state levels of Nb22-activated WT LRRK2, resembling the pronounced CYLD-dependent increase previously observed for the pathogenic Y1699C variant (Eckert *et al*., 2026). In contrast, Nb17-bound WT LRRK2 exhibited higher basal steady-state levels than Nb22-activated LRRK2, and its abundance was not significantly further increased by CYLD co-expression (Figure 5A; Supplementary Figure 10A). Thus, despite identical CYLD co-expression conditions, the effect of CYLD on LRRK2 abundance differed between the two nanobody-defined states. Interestingly, we found a reduced relative pS935 levels for both nanobodies in presence of CYLD (Supplementary Figure 10B). However, CYLD increased Nb22-bound LRRK2 abundance without a proportional increase in pT73-Rab10 relative to total Rab10, resulting in reduced Rab10-directed signaling output per unit LRRK2 (Supplementary Figure 10C, D). In Nb17-expressing cells, pRab10 remained below the limit of detection, consistent with inhibition of LRRK2 kinase activity by Nb17.

**Figure 5:**
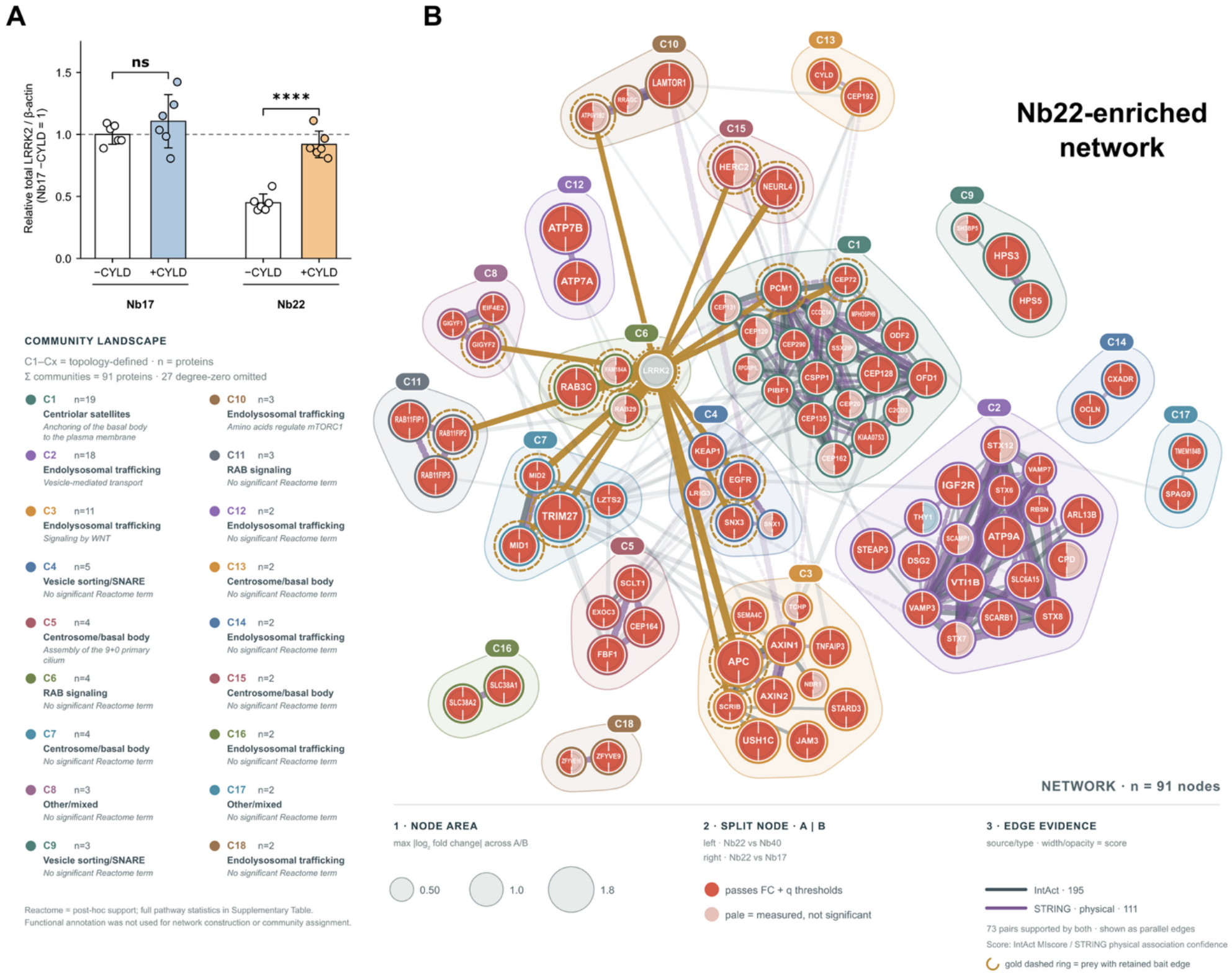
Functional dissection of the Nb22-associated LRRK2 proximity network. (A) Effect of CYLD co-expression on steady-state levels of nanobody-bound WT LRRK2. HEK293T cells expressing WT LRRK2 together with Nb17 or Nb22 were analyzed in the absence or presence of co-expressed CYLD. LRRK2 abundance was quantified by capillary Western analysis using the 24D8 antibody and normalized to β-actin. Values are shown relative to the mean of the Nb17 −CYLD condition. Bars represent mean ± SD and individual data points represent biological replicates (N = 6). Statistical significance was assessed separately for the effect of CYLD within the Nb17- and Nb22-bound LRRK2 conditions using two-sided unpaired Student’s t-tests. ns, not significant; ****P < 0.0001. (B) Consensus protein-association network integrating Nb22 vs Nb40 and Nb22 vs Nb17. Proteins were included if they were identified in both and met the active fold-change and q-value criteria in at least one of the two comparisons. Nodes are displayed as dual-effect nodes, with the left and right halves representing Nb22 vs Nb40 and Nb22 vs Nb17, respectively. Node area scales with the strongest absolute log2 fold change across the two comparisons. Node fill indicates the direction and magnitude of differential abundance, with orange denoting enrichment with Nb22. Fully saturated node halves indicate comparisons that meet the active fold-change and q-value criteria, whereas paler halves indicate measured effects that do not meet these criteria. Colored node rings and hulls denote connectivity-defined subnetworks identified by weighted Leiden-style community decomposition. Solid dark edges denote IntAct interactions, whereas dashed purple edges denote STRING physical associations. Edge width and opacity scale with IntAct MIscore (cut-off: 0.35) and STRING physical-association confidence (cut-off: 0.40), respectively. Protein pairs supported independently by both databases are displayed as parallel edges. The LRRK2 bait is highlighted. Communities were functionally annotated post hoc, including with Reactome pathway enrichment. The network comprises 91 connected nodes partitioned into 18 communities (C1–C18); 27 proteins without network connections are not shown.

However, the Nb22-associated interactome also displayed features that were absent or less evident in the inhibitor-stabilized state. Among the most strongly enriched proteins were multiple components of the endolysosomal system, including the Ragulator complex component LAMTOR1. In addition, several RAB GTPases were significantly enriched in the Nb22 proximity proteome, including the established LRRK2 upstream regulator RAB29 (Figure 3C, 4C). While RAB10 was robustly detected as an LRRK2-proximal protein in the biotin-labeling datasets (Supplementary Figure 3), only a trend toward enrichment in the Nb22 (activated LRRK2) condition was observed in the comparative analyses. In contrast, RAB3C, a proposed LRRK2 substrate (Nirujogi *et al*, 2021; Steger *et al*, 2017), and RAB33B were significantly enriched in the Nb22-associated protein network.

In addition to RAB GTPases, the activated LRRK2 conformation selectively enriched several established components of the LRRK2 signaling network, including the RAB effectors SPAG9 (JIP4), RILPL1, and members of the MICAL/MICALL family (Bonet-Ponce *et al*., 2020; Dhekne *et al*, 2018; Ordóñez *et al*., 2019; Steger *et al*., 2016). Notably, the E3 ubiquitin ligase MID2 (TRIM1), which has been shown to regulate LRRK2 localization, ubiquitination, and degradation, was also enriched in the Nb22-associated proximity proteome (Stormo *et al*, 2022). Together, these findings indicate that stabilization of the active LRRK2 conformation promotes the recruitment of both upstream regulatory proteins and downstream signaling effectors into its molecular neighborhood.

To examine whether proteins enriched in the Nb22 proximity proteome form an interconnected molecular network, we integrated the Nb22-versus-Nb40 Nb22-versus-Nb17 datasets with protein-association information from STRING and IntAct (Figure 5B). Connectivity-based analysis revealed several densely interconnected subnetworks linked by a smaller number of bridging proteins. Importantly, subnetwork assignment was based exclusively on network topology and was independent of functional annotation. The resulting network recapitulated the major biological signatures identified by the differential proteomic and GO analyses, with prominent interconnected groups containing centrosomal and centriolar-satellite proteins, including PCM1, OFD1, SSX2IP and CEP131, together with RAB-associated and endolysosomal proteins such as RAB29, RAB3C, RAB33B, LAMTOR1, RILPL1 and SPAG9. Thus, rather than representing isolated changes in protein proximity, Nb22-enriched proteins form an organized association network linking centrosomal/ciliary and membrane-trafficking components of the LRRK2 molecular environment.

Overall, the Nb22-versus-Nb40 and Nb22-versus-Nb17 comparisons yielded highly similar enrichment profiles, demonstrating robust remodeling of the LRRK2 molecular neighborhood by the activating nanobody. The Nb22-associated proximity proteome combined features of the centrosome/cilium-associated environment observed upon type I inhibitor treatment with the RAB/endolysosomal environment associated with RAB29-dependent LRRK2 signaling (Supplementary Figure 11). Thus, the Nb22-stabilized conformation appears to define a molecular neighborhood that integrates components of previously distinct LRRK2 signaling environments. A schematic representation of these major modules is shown in Figure 6A.

**Figure 6:**
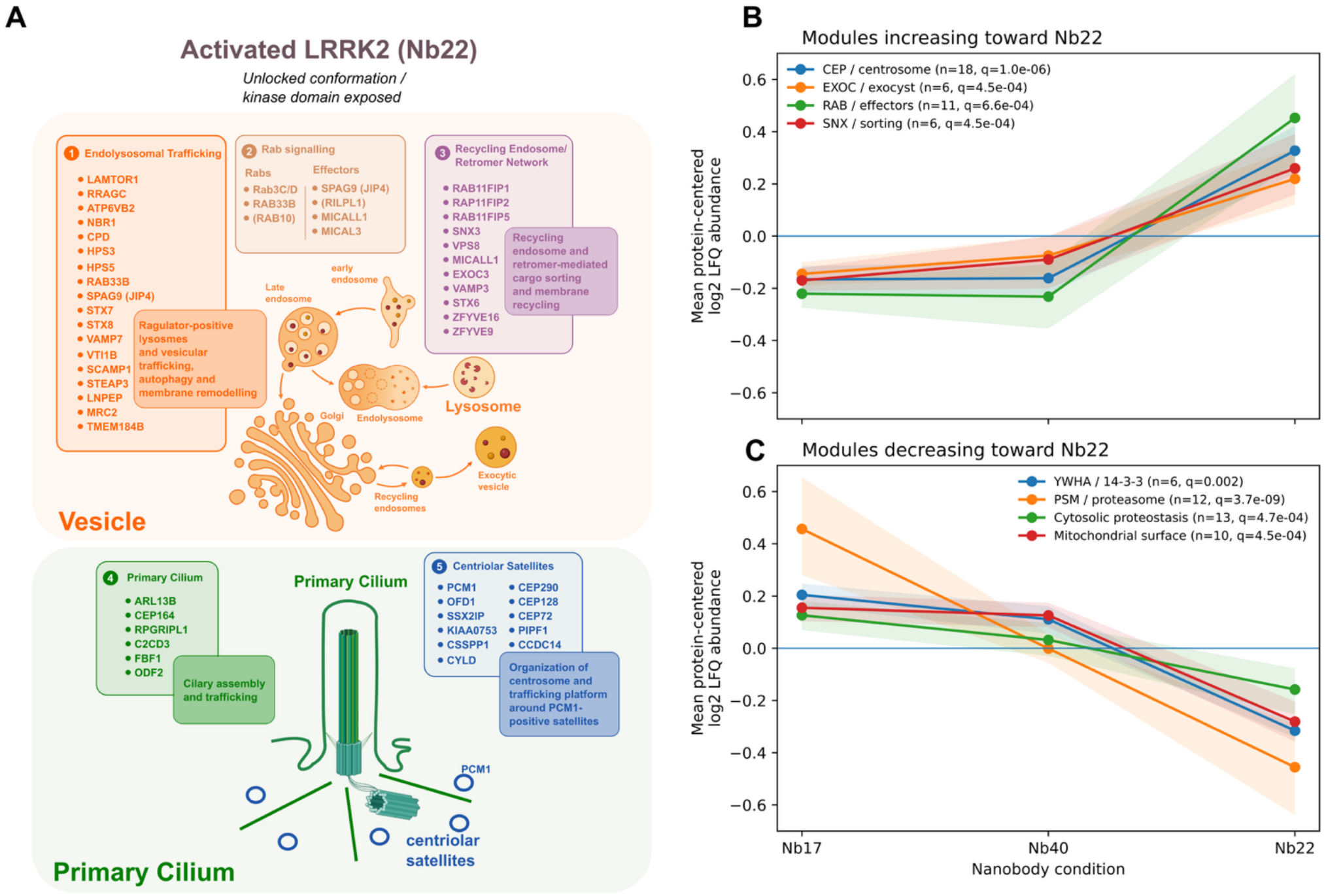
Activation-state-specific remodeling of the LRRK2 proximity network. (A) Schematic summary of the proximity-proteomic landscape associated with the activated, “unlocked” conformation of LRRK2 induced by the activating WD40-binding nanobody Nb22. Key proteins significantly enriched in the Nb22 proximity proteome relative to the largely neutral WD40-binding nanobody Nb40 or the inhibitory Nb17 were grouped into functionally related modules based on Gene Ontology annotations and literature curation. Five major cellular compartments and pathways emerge from the activated LRRK2 proximity network. (B, C) Ordered remodeling of protein groups across the nanobody-defined LRRK2 states. Protein families associated with the Nb22-versus-Nb17 discovery contrast were identified by data-driven family enrichment and consolidated into a fixed set of nonredundant modules before three-state trend analysis. Exploratory cytosolic-proteostasis and mitochondrial-surface protein sets were analyzed using the same framework. Mean protein-centered log₂ LFQ abundances are shown across Nb17, Nb40, and Nb22. Lines represent the mean of all quantified proteins within each group, and shaded areas indicate bootstrap 95% confidence intervals calculated from 10,000 resamplings of proteins. Module size (n) and Benjamini–Hochberg-adjusted P values (q) from Page’s ordered trend test are indicated in the legends. Centrosomal CEP, EXOC/exocyst, RAB/RAB-associated, and SNX/sorting-nexin modules progressively increased toward Nb22 (B), whereas PSM/proteasomal, cytosolic-proteostasis, mitochondrial-surface, and YWHA/14-3-3 groups showed reciprocal decreases (C). Independent +/−biotin comparisons were used as an orthogonal assessment of proximity-labeling specificity but not as an absolute inclusion criterion. YWHA/14-3-3 was retained as a predefined special case because high nonspecific recovery can reduce apparent biotin-dependent enrichment, while its trajectory is independently consistent with the corresponding changes in LRRK2 S935 phosphorylation. Representative anchor proteins were not used to calculate module trajectories. Bootstrap confidence intervals reflect variation across proteins within each group and do not represent biological-replicate confidence intervals.

### Data-driven analysis reveals ordered remodeling of LRRK2 molecular neighborhoods

To determine whether differences observed in the pairwise Nb22 comparisons reflected isolated protein-level changes or coordinated remodeling of the LRRK2 molecular environment, we performed a data-driven analysis of the processed Nb22-versus-Nb17 and Nb22-versus-Nb40 proximity-proteomic datasets. The Nb22-versus-Nb17 comparison was used as the discovery contrast. Proteins were ranked according to the direction and significance of differential abundance, and protein families overrepresented among the most strongly Nb22-enriched or Nb22-depleted proteins were identified without prior selection of individual anchor proteins. The analysis recovered coherent protein families with reciprocal behavior. Families enriched toward Nb22 included centrosomal CEP proteins, EXOC/exocyst components, RAB proteins and RAB-associated factors, and sorting nexins (SNX), whereas depleted families included proteasomal PSM components and YWHA/14-3-3 proteins. Following removal of redundant assignments, the retained data-derived families were fixed before subsequent analysis across the three nanobody conditions. In addition, two exploratory groups were evaluated independently of the family-discovery procedure: a cytosolic-proteostasis set comprising chaperone, redox, and peptidyl-prolyl isomerase proteins, and a mitochondrial-surface set comprising cytosol-accessible mitochondrial proteins selected using localization annotation together with biotin-dependent proximity support. Independent +/−biotin comparisons were used as an orthogonal assessment of proximity-labeling specificity rather than as an absolute exclusion criterion.

We next tested whether these protein groups exhibited ordered behavior across the inhibitory Nb17, largely neutral Nb40, and activating Nb22 conditions. A pronounced reciprocal organization emerged (Figure 6B and C). The CEP/centrosomal module progressively increased from Nb17 through Nb40 to Nb22 (n = 18, FDR = 1.0 × 10⁻⁶). Similar ordered increases were observed for EXOC/exocyst and SNX/sorting-nexin proteins (both n = 6, FDR = 4.5 × 10⁻⁴) and the RAB/RAB-associated module (n = 11, FDR = 6.6 × 10⁻⁴). Five of six EXOC and five of six SNX proteins individually followed the strict Nb17 < Nb40 < Nb22 ordering.

Conversely, the PSM/proteasomal module showed the strongest ordered decrease (n = 12, FDR = 3.7 × 10⁻⁹), with all 12 proteins following the strict Nb17 > Nb40 > Nb22 ordering. The exploratory cytosolic-proteostasis and mitochondrial-surface sets likewise decreased toward Nb22 (n = 13, FDR = 4.7 × 10⁻⁴ and n = 10, FDR = 4.5 × 10⁻⁴, respectively). The mitochondrial-surface trajectory was supported by biotin-dependent enrichment of its constituent proteins, consistent with state-dependent changes in LRRK2 proximity to the cytosolic face of mitochondria. YWHA/14-3-3 proteins also decreased coherently (n = 6, FDR = 1.6 × 10⁻³). Although YWHA proteins showed limited positive enrichment in the +/−biotin specificity comparisons, they were retained as a distinct regulatory family because nonspecific recovery can reduce apparent biotin-dependent enrichment and their state-dependent trajectory was independently consistent with the opposing effects of Nb17 and Nb22 on LRRK2 S935 phosphorylation and its phosphorylation-dependent association with 14-3-3 proteins.

Together, the frequent intermediate position of Nb40 and reciprocal behavior of multiple protein systems indicate progressive remodeling of the LRRK2 molecular environment across the nanobody-defined states. The data are consistent with redistribution from cytosolic-proteostasis-, mitochondrial-surface-, and 14-3-3-associated environments toward centrosomal and membrane-trafficking neighborhoods in the Nb22-associated activated state.

## Discussion

Recent structural studies have transformed our understanding of LRRK2 from a catalytic enzyme into a highly dynamic multidomain signaling protein whose activity is intimately linked to its conformational state (Deniston *et al*., 2020; Myasnikov *et al*., 2021; Schmidt *et al*., 2021; Villagran Suarez *et al*., 2026; Zhu *et al*., 2023). While cryo-EM and biochemical analyses have revealed the existence of autoinhibited and active conformations, the cellular consequences of these structural transitions remain incompletely understood. In the present study, we combined nanobody-mediated stabilization of distinct LRRK2 states with proximity proteomics to investigate how perturbation of the conformational landscape influences the local molecular environment of the kinase. Our findings reveal extensive remodeling of the LRRK2 proximity proteome upon stabilization of an activated state and support a model in which conformational transitions contribute to the spatial organization of LRRK2 signaling networks.

Accumulating structural and biochemical evidence suggests that the WD40 domain participates in intramolecular interactions that contribute to the regulation of LRRK2 activity (Myasnikov *et al*., 2021; Villagran Suarez *et al*., 2026). Consistent with this notion, both nanobodies and DARPins targeting the WD40 domain have been shown to modulate LRRK2 function, indicating that this region acts as an important regulatory node within the protein (Dederer *et al*, 2024; Singh *et al*., 2022). Our findings extend these observations by demonstrating that perturbation of the WD40 domain is accompanied by extensive remodeling of the LRRK2 proximity proteome, suggesting that WD40-mediated conformational transitions not only regulate kinase activity but also influence the organization of LRRK2-associated molecular environments.

While further experimental validation using cryo-EM is required, the structural models generated in this study provide a plausible mechanistic explanation for the activating properties of Nb22. The predicted binding site is located adjacent to the recently described hinge helix, a structural element implicated in maintaining the autoinhibited conformation of LRRK2 (Myasnikov *et al*., 2021). Superposition of the Nb22-bound model onto the autoinhibited cryo-EM structure suggests steric incompatibility with the folded arrangement of the N-terminal domains, implying that nanobody binding may destabilize long-range intramolecular contacts that maintain the inactive state. These models are thus consistent with the observed increase in RAB10 phosphorylation, the loss of S935 phosphorylation, and recent structural analyses of active monomeric LRRK2. Furthermore, the LRRK2 missense variant G2385R, a common Parkinson’s disease risk variant in the Han Chinese population, is located within the predicted Nb22 binding epitope. Notably, this variant has been reported to increase LRRK2 kinase activity by twofold, as evidenced by enhanced RAB10 phosphorylation in cells (Kalogeropulou *et al*., 2022; Nguyen *et al*., 2026; Tezuka *et al*., 2022). Our data recapitulate the elevated basal activity of G2385R and further show that Nb22 produces only a modest additional increase in RAB10 phosphorylation, which remains substantially below the level observed for Nb22-bound wild-type LRRK2. Together with the absence of Nb22-induced S935 dephosphorylation in the G2385R background, these findings are consistent with impaired productive engagement of Nb22 by the G2385R variant and support the proposed binding model. In contrast, the predicted binding mode of Nb40 is fully compatible with the autoinhibited structure and provides a structural rationale for its largely neutral behavior. Together, these observations support a model in which Nb22 promotes activation by destabilizing autoinhibitory interactions involving the WD40 domain and the N-terminal region of LRRK2.

The most striking observation emerging from our proteomic analyses is the extensive overlap between the Nb22-associated interactome and the molecular environment previously observed following treatment with type I kinase inhibitors. At first glance this appears paradoxical, as Nb22 activates LRRK2 whereas MLi-2 suppresses catalytic activity. Although type I inhibitors inhibit phosphotransferase activity, they stabilize the closed, active-like DYG-in conformation of the LRRK2 kinase domain, whereas type II inhibitors stabilize the open, inactive DYG-out state (Alessi & Pfeffer, 2024; Zhu *et al*, 2024). The substantial overlap between both datasets therefore suggests that many aspects of LRRK2-associated molecular organization are determined by conformational state rather than kinase activity alone. Particularly notable was the shared enrichment of centriolar satellite proteins including PCM1, OFD1, CEP131 and SSX2IP. Given the central role of centriolar satellites in organizing centrosomal and ciliary protein networks, these findings further support the notion that specific LRRK2 conformations preferentially engage centrosome-associated molecular assemblies. Recent work further demonstrates that centriolar satellites are dynamically organized PCM1-based structures in which client proteins are recruited hierarchically and remodeled in a context-dependent manner (Begar *et al*, 2026). Moreover, the centrosome– cilium–satellite axis is increasingly implicated in neurodegenerative disease, while disruption of centrosomal and ciliary homeostasis is well established in the context of pathogenic LRRK2 signaling (Sahin & Firat-Karalar, 2026). Despite this overlap, the activated Nb22-associated interactome does not fully recapitulate the inhibitor-stabilized state. Several proteins preferentially enriched in the Nb22 datasets have previously been linked to membrane trafficking and endolysosomal biology. Among these were the Ragulator component LAMTOR1, the established upstream regulator RAB29, the proposed LRRK2 substrates RAB3C/D and RAB33B, and SPAG9/JIP4, a component implicated in LRRK2-dependent lysosomal tubulation pathways (Bonet-Ponce *et al*., 2020). Collectively, these observations suggest that activation through WD40-directed nanobodies promotes recruitment of signaling assemblies associated with endolysosomal membranes and RAB-dependent trafficking pathways. The simultaneous enrichment of centrosomal and endolysosomal components is particularly intriguing and suggests that the Nb22-stabilized state cannot be explained by a simple recapitulation of previously characterized signaling environments.

RAB3C was among the proteins most prominently enriched in the Nb22-associated LRRK2 proximity proteome. Its increased proximity to the Nb22-stabilized, unlocked LRRK2 state is particularly notable because RAB3 family proteins are key regulators of synaptic vesicle trafficking, and disruption of RAB3 family function has been associated with neurological disease (He *et al*, 2025; Hengel *et al*, 2025; Schluter *et al*, 2004). In addition, accumulating evidence links LRRK2 to RAB3-dependent trafficking pathways, and RAB3C has emerged as a putative LRRK2 substrate (Pfeffer, 2023; Twellsieck & Boecker, 2025). Consistent with this, LRRK2 regulates presynaptic vesicle homeostasis, with depletion of LRRK2 altering synaptic vesicle distribution, recycling dynamics, and neurotransmission (Piccoli *et al*, 2011). Furthermore, GST pull-down experiments identified an interaction between RAB3A and the isolated LRRK2 WD40 domain, providing biochemical support for a functional association between LRRK2 and RAB3-mediated vesicle trafficking (Piccoli *et al*, 2014). More recently, pathogenic LRRK2 was shown to impair anterograde transport of synaptic vesicle precursors through phosphorylation of the RAB3 family member RAB3A, thereby disrupting its interactions with key trafficking regulators (Dou *et al*, 2024). In addition, Drosophila studies have provided further evidence linking LRRK2 to RAB3-dependent vesicle biology (Islam *et al*, 2016). Although our proximity labeling data do not demonstrate a direct functional interaction, the selective enrichment of RAB3C supports the hypothesis that stabilization of the activated LRRK2 conformation promotes association with molecular machinery involved in RAB3-dependent vesicle transport. Together, these findings support our broader model in which distinct LRRK2 conformational states organize specialized molecular neighborhoods, linking activation-state–dependent interactome remodeling to synaptic vesicle trafficking pathways that are central to neuronal function and are disrupted in PD.

One of the most notable findings of the present study is that the Nb22-associated proximity proteome combines features previously observed in two apparently distinct LRRK2 signaling states: the centrosome-associated interactome induced by type I kinase inhibitors and the membrane-associated interactome associated with RAB29-mediated activation (Eckert *et al*., 2026). This observation suggests that Nb22 does not merely mimic either perturbation but instead accesses a distinct region of the LRRK2 conformational landscape. Such an interpretation is consistent with recent structural studies indicating that LRRK2 occupies a continuum of conformational states rather than a simple binary active/inactive equilibrium (Myasnikov *et al*., 2021; Park *et al*, 2026; Villagran Suarez *et al*., 2026; Weng *et al*., 2023). Within this framework, distinct perturbations may stabilize overlapping but non-identical structural states, resulting in partially shared molecular neighborhoods. The ability of Nb22 to capture features of both inhibitor-induced and RAB29-induced signaling programs therefore provides experimental evidence that discrete conformational states are associated with distinct interaction networks. We recently identified CYLD as a candidate LRRK2 effector through integrated proximity-proteomic and co-evolutionary analyses and found particularly strong evolutionary coupling to the ROC-COR region of LRRK2, including the pathogenic hotspot residue Y1699 (Eckert *et al*., 2026). In the present study, CYLD was again enriched in the activated LRRK2 proximity proteome and selectively increased the steady-state abundance of Nb22-activated WT LRRK2, but not Nb17-bound WT LRRK2. This differential response under identical CYLD co-expression conditions supports a dependence of the effect on the nanobody-defined LRRK2 state rather than a uniform consequence of CYLD overexpression. Together with the preferential CYLD-dependent increase previously observed for Y1699C LRRK2, these findings provide orthogonal functional support for a conformational-state-dependent relationship between CYLD and LRRK2. Notably, the CYLD-dependent increase in Nb22-bound LRRK2 abundance was not accompanied by a proportional increase in Rab10 phosphorylation, resulting in reduced Rab10-directed signaling output per unit LRRK2. This uncoupling suggests that CYLD increases the abundance of an LRRK2 pool that does not contribute proportionally to downstream Rab10 signaling. Rather than simply enhancing kinase output, CYLD may therefore influence the stability, turnover, or subcellular partitioning of specific LRRK2 states. Whether the additional CYLD-responsive LRRK2 pool differs in intrinsic catalytic activity, substrate accessibility, or localization remains to be determined.

This relationship may also connect LRRK2 conformational regulation to the centriolar-satellite network. CYLD was enriched together with core centriolar-satellite components, including PCM1, OFD1, CEP131 and SSX2IP, and has previously been shown to regulate centriolar-satellite proteostasis by counteracting the E3 ubiquitin ligase MIB1 (Douanne *et al*., 2019). Moreover, recent work demonstrated a role for CYLD-mediated K63 deubiquitination in neuronal proteostasis, synaptic function and autophagy/autolysosomal organization (Sotiriou *et al*, 2026). The Nb22 proximity proteome additionally contained MID2/TRIM1, an E3 ubiquitin ligase implicated in LRRK2 ubiquitination, localization and turnover in relation to its S910/S935 phosphorylation state (Stormo *et al*., 2022). Thus, the coordinated enrichment of CYLD, MID2 and centriolar-satellite proteins raises the possibility that reversible ubiquitination contributes to the organization or turnover of active-state LRRK2 signaling environments. Given the established role of centriolar satellites in centrosomal protein quality control and aggresome pathways, the Nb22-associated satellite environment may therefore also reflect state-dependent regulation of LRRK2 proteostasis (Prosser *et al*, 2022).

The data-driven analysis further supports coordinated remodeling of the LRRK2 molecular environment across nanobody-defined states. Centrosomal and membrane-trafficking families progressively increased from Nb17 through Nb40 to Nb22, in parallel with increasing RAB10 phosphorylation, whereas proteasomal, cytosolic-proteostasis, and mitochondrial-surface signatures showed the reciprocal behavior. The mitochondrial-surface trajectory is particularly notable because it was supported by proteins accessible from the cytosolic face of mitochondria, arguing against an interpretation based solely on internal mitochondrial proteins. YWHA/14-3-3 proteins represented a distinct regulatory signature whose decrease is consistent with the strong reduction in LRRK2 S935 phosphorylation in the Nb22 state. Together, these reciprocal trajectories are consistent with state-dependent redistribution of LRRK2 from a more cytosolic and mitochondrial-associated molecular environment toward vesicular and pericentrosomal neighborhoods.

More broadly, our findings support a model in which LRRK2 functions not only as a kinase but also as a conformationally regulated signaling platform whose local molecular environment changes with its structural state. In this view, activation involves not only altered substrate phosphorylation but also redistribution of local molecular neighborhood across centrosomal, ciliary, vesicular and endolysosomal pathways. Such a mechanism may help explain how pathogenic mutations located in distinct regions of LRRK2 converge on common cellular phenotypes despite exerting diverse biochemical effects. Importantly, these observations further suggest that conformational state itself may represent a therapeutically relevant layer of LRRK2 regulation.

Several limitations of the present study should be acknowledged. Proximity labeling identifies proteins within the local environment of LRRK2 but does not distinguish direct interactions from indirect spatial associations. Furthermore, although the nanobodies employed here provide unique tools to manipulate the conformational landscape of LRRK2, the precise structural states stabilized in living cells remain incompletely defined. Likewise, the structural interpretations presented here are based primarily on AlphaFold3 predictions and therefore remain hypothesis-generating. Future studies combining nanobody stabilization with cryo-EM and advanced cellular imaging approaches will be required to define more precisely the structural basis of the observed molecular neighborhoods.

In summary, our findings show that nanobody-mediated perturbation of the LRRK2 conformational landscape is accompanied by extensive remodeling of its local molecular environment. The activated Nb22-stabilized state combines a centrosomal and ciliary signature shared with type I inhibitor-treated LRRK2 with an endolysosomal and Rab-associated module characteristic of membrane-associated LRRK2 signaling. Together, these findings support the concept that distinct LRRK2 conformational states are associated with distinct molecular neighborhoods. More broadly, this study establishes conformation-selective nanobodies coupled with proximity proteomics as a strategy for linking structural states of dynamic multidomain proteins to their cellular signaling environments.

## Methods

### Cloning of the expression constructs

The HA-miniTurbo-LRRK2 (mT-LRRK2) construct was previously described (Eckert *et al*., 2026). For cell-based LRRK2 kinase assays, N-terminally Strep-FLAG tagged WT LRRK2 was used and pathogenic variants R1441G and G2385R were introduced by site-directed mutagenesis as previously described (Gilsbach *et al*, 2024). The nanobodies C-terminally tagged with eGFP were previously described in (Singh *et al*., 2022).

### Cell culture and biotin labeling

Cell culture conditions and proximity labeling procedures were performed essentially as described previously (Eckert *et al*., 2026). Briefly, HEK293T cells (CVCL_0063) were maintained in Dulbecco’s modified Eagle medium (DMEM) supplemented with 10% fetal bovine serum and 0.5% penicillin/streptomycin at 37°C and 5% CO₂. The genetic characteristics of this cell line were determined previously (Gilsbach *et al*., 2024). Cells were co-transfected with mT-LRRK2 and eGFP-tagged nanobody constructs using polyethyleneimine (PEI) 25 kDa (Polysciences) as described in (Ben-Zimra *et al*, 2002). For proximity labeling experiments, cells were seeded in 14 cm dishes and transfected with plasmids encoding HA-miniTurbo-LRRK2 and the respective nanobody at a ratio of 70:30. Per biological replicate and condition, one 14 cm dish was used. Proximity labeling was initiated by addition of biotin to a final concentration of 50 µM for 2 h at 37°C. Following labeling, cells were washed with ice-cold PBS and harvested for protein extraction. For the replication of the inhibitor experiments, initially published in Eckert *et al*., 2026 (MLi-2 vs GZD-824 vs DMSO), a detailed protocol was previously published on Zenodo (https://doi.org/10.5281/zenodo.17045153).

### Enrichment of biotinylated proteins

Protein extraction and affinity enrichment of biotinylated proteins was performed as previously described (Eckert *et al*., 2026) with minor modifications. Briefly, cells were lysed in 1 mL lysis buffer [TBS (30 mM Tris-HCl, pH 7.4, and 150 mM NaCl) supplemented with 1% NP-40 substitute (Merck), protease (Roche) and phosphatase inhibitors (PhosSTOP, Roche)] per 14 cm dish. Lysates were incubated for 20 min at 4°C with gentle mixing and clarified by centrifugation at 1,000 × g for 10 min at 4°C. The resulting supernatants were adjusted to a final concentration of 1% SDS and incubated for an additional 15 min at 4°C with gentle mixing. Following centrifugation at 20,000 × g for 10 min at room temperature, cleared lysates were transferred to fresh tubes and protein concentrations were determined using a BCA assay.

For affinity enrichment, equal amounts of total protein were incubated overnight at 4°C with MagStrep® Strep-Tactin® beads (#2-1613-002, IBA Lifesciences) under continuous rotation. Prior to use, beads were washed three times with TBS containing 0.1% NP-40 substitute. Following overnight incubation, beads were collected using a magnetic rack and washed four times with TBS containing 0.1% NP-40 substitute to remove non-specifically bound proteins.

Bound proteins were eluted with elution buffer BXT (2-1042-025, IBA Lifesciences). Eluted proteins were precipitated by chloroform/methanol precipitation as described in (Gloeckner *et al*, 2009) and subsequently processed for downstream proteomic analysis by in-solution digestion and liquid chromatography–tandem mass spectrometry (LC-MS/MS) as described below.

### Mass spectrometry

The protein samples were subjected to proteolytic cleavage as described earlier before the resulting peptides were analyzed by LC-MS (Gloeckner *et al*., 2009). Peptides were separated using a nanoElute 2 UHPLC system (Bruker) coupled to a timsTOF Ultra 2 mass spectrometer (Bruker). A trap-column configuration comprising a Thermo Trap Cartridge (5 mm) and a PepSep Twenty-five Series analytical column (75 μm inner diameter, 1.5 μm particle size) was used, with the analytical column maintained at 50°C.

Peptides were loaded using an ultra-low-volume pickup injection (four pickup cycles plus 2 μL) and separated at a flow rate of 0.30 μL min⁻¹ using mobile phase A (0.1% formic acid in water) and mobile phase B (0.1% formic acid in 99.9% acetonitrile). The gradient was 5–25% B over 24 min, 25–35% B over 6 min, increased to 95% B by 34 min, held until 38 min, and re-equilibrated at 5% B until 40 min.

Mass spectrometric data were acquired in DIA-PASEF mode (Meier *et al*, 2020) using a CaptiveSpray nanoelectrospray ion source operated in positive-ion mode with a spray voltage of 1.6 kV. TIMS separation was performed over an ion mobility range of 0.60–1.40 Vs cm⁻². MS spectra were acquired over an m/z range of 100– 1700, while DIA-PASEF windows covered precursor ions between m/z 300 and 1200. The acquisition method comprised 12 DIA-PASEF scans per cycle with an accumulation time of 2 ms, a ramp time of 100 ms, and a total cycle time of approximately 1.38 s. Fragmentation was performed by collision-induced dissociation (CID) using ion mobility-dependent collision energies, and data were acquired in high-sensitivity mode.

### Mass spectrometry data analysis

Raw DIA-PASEF data were analyzed using DIA-NN (v2.6.0) (Demichev *et al*, 2020). Searches were performed against a custom FASTA database comprising the reviewed human UniProt proteome (release 2025_05) and expressed nanobody sequences. DIA-NN was run with FASTA search enabled, tryptic digestion specificity assuming tryptic cleavage after lysine and arginine, up to one missed cleavage, N-terminal methionine excision enabled, and cysteine carbamidomethylation as a fixed modification. Peptides with lengths between 7 and 45 amino acids, precursor charge states from 2 to 4, precursor m/z values between 300 and 1450, and fragment m/z values between 200 and 2000 were considered. MS1 and MS2 mass accuracies were fixed at 15 ppm. Match-between-runs was enabled, and output was filtered at 5% FDR. Precursor-and protein-level expression matrices were exported for downstream statistical analysis.

For downstream analyses, nanobody sequences and keratins were excluded. Statistical analyses were performed using an in-house Python workflow, and graphical representations were generated using the Python packages Matplotlib and Seaborn.

Protein groups were retained if quantitative values were present in all but one sample in at least one experimental condition. Missing values were imputed using an MNAR algorithm adapted from *msImpute* (Lazar *et al*, 2016). Differential protein abundance was assessed using a limma-like python implementation of moderated *t*-statistics (Smyth, 2004). To account for uncertainty introduced by missing-value imputation, differential abundance testing was performed across 100 independently imputed datasets and results were combined according to Rubin’s rules (Rubin, 1987). Volcano plots display the log2 fold change against the –log10 adjusted *P* value derived from the Rubin-pooled analysis, with statistical significance determined using the Benjamini–Hochberg false discovery rate (FDR). In addition to statistical significance (FDR-adjusted q-value), effect size thresholds were applied to identify enriched proteins. For the +biotin versus −biotin comparisons, proteins with a log₂ fold change (log₂FC) ≥ 1.0 were considered enriched. For comparisons between nanobody conditions and inhibitor treatments, a threshold of log₂FC ≥ 0.5 was applied. Bar charts of selected proteins show mean normalized protein intensities ± SD across replicates, and the reported q-values correspond to the Benjamini–Hochberg-adjusted results from the limma-like analysis.

### Capillary Western Blots

Capillary Western blot analyses were performed using the Jess automated capillary electrophoresis system (Bio-Techne) as previously described (Eckert *et al*., 2026), with minor modifications. Protein separation was carried out using a 25-capillary fluorescence separation cartridge (12–230 kDa; SM-FL004-1) together with the Jess Capillary Western kit (SM-W003; Bio-Techne), according to the manufacturer’s instructions. Cell lysates expressing LRRK2 variants together with GFP-tagged nanobodies or an empty vector control (70:30 ratio) were loaded at 1.25 µg total protein per capillary. The capillaries were blocked with milk-free Antibody Diluent, Bio-Techne, 043-524.

For LRRK2 analyses, phosphorylated LRRK2 (S935) was detected using a rabbit monoclonal anti-pS935-LRRK2 antibody (Abcam, ab133450; 1:50 in milk-free Antibody Diluent). Total LRRK2 was detected using an in-house generated rat monoclonal antibody (clone 24D8; 1:25), followed by a rabbit anti-rat DyLight-633 secondary antibody (Bio-Techne, NBP1-76225; 1:50 in milk-free Antibody Diluent).

Rab10 phosphorylation was assessed using a rabbit monoclonal anti-pT73-Rab10 antibody (Abcam, ab230261; 1:40 in milk-free Antibody Diluent), while total Rab10 was detected using a rabbit monoclonal anti-Rab10 antibody (Abcam, ab181367; 1:20 in milk-free Antibody Diluent). Both antibodies were detected using an HRP-conjugated anti-rabbit secondary antibody (Bio-Techne, 042-206).

Expression of GFP-tagged nanobodies was determined using an anti-GFP antibody (rat monoclonal antibody, generated in-house, clone 3H9, 1:20 in milk-free Antibody Diluent). β-actin was used as a loading control and detected using a mouse anti-β-actin antibody (R&D Systems, MAB8929; 1:500 in milk-free Antibody Diluent), followed by an anti-mouse near-infrared secondary antibody (Bio-Techne, 043-821).

Phosphorylated and total LRRK2 were analyzed on one Jess array, phosphorylated and total Rab10 on a second array, and LRRK2 together with β-actin on a third array. Expression levels of the GFP-Nb constructs were tested separately. Sequential probing was performed by stripping capillaries with RePlex buffer (RePlex module, Bio-Techne, RP-001) between detection rounds.

Electropherograms were analyzed using Compass for SW software (version 7.1.0; Bio-Techne). Quantitative analyses were performed using custom Python scripts, and graphical representations were generated in Python using Matplotlib. Because total LRRK2 expression differed substantially between LRRK2 variants and nanobody conditions, RAB10 phosphorylation was first normalized to total RAB10 and subsequently to β-actin-normalized total LRRK2 abundance. This additional normalization estimated RAB10 phosphorylation relative to the amount of expressed LRRK2. For each experimental series, values were normalized to the corresponding WT-EV mean. Positive normalized values were log₂-transformed before statistical testing. Within each genotype, two-sided Dunnett tests were used to compare Nb17 and Nb22 with the EV control, with adjustment for the two comparisons. WT observations from the WT/R1441G and WT/G2385R experimental series were combined only after normalization to the corresponding within-series WT-EV mean.

### CYLD co-expression and LRRK2 quantification

CYLD co-expression experiments were performed essentially as described previously (Eckert *et al*., 2026). HEK293T cells were co-transfected with SF-TAP-tagged WT LRRK2, Nb17 or Nb22, and either CYLD or the corresponding empty-vector control at a plasmid ratio of 3:1:6, respectively. Cells were harvested 48 h after transfection and lysed as described above. LRRK2 steady-state levels were quantified using the JESS capillary Western system (Bio-Techne) with the 24D8 LRRK2 antibody and normalized to β-actin. Rab10 phosphorylation was quantified on a separate Jess array as pT73-Rab10 relative to total Rab10 using the antibodies described above. For the Nb22 condition, the pT73-Rab10/total Rab10 ratio was additionally normalized to the corresponding β-actin-normalized LRRK2 abundance from the same biological replicate to estimate Rab10-directed signaling output per unit LRRK2. LRRK2-normalized values were expressed relative to the mean of the Nb22 −CYLD condition. In Nb17-expressing cells, pT73-Rab10 was below the limit of detection and was therefore not subjected to quantitative or statistical analysis. Statistical significance of CYLD-dependent changes in LRRK2 abundance was assessed separately for Nb17- and Nb22-bound LRRK2 using two-sided unpaired Student’s t-tests. For the Nb22 Rab10 analyses, −CYLD and +CYLD measurements derived from matched biological experiments were compared using two-sided paired Student’s t-tests. A p value < 0.05 was considered statistically significant.

### Gene Ontology enrichment analysis

Functional enrichment analysis was performed on proteins identified as significantly enriched in the respective pairwise comparisons. Gene Ontology (GO) Biological Process enrichment was carried out using the Python package GSEApy, employing the Enrichr database with the Gene Ontology (GO) Biological Process (2023 release) gene set and the human reference annotation. Only proteins classified as significantly enriched by the statistical analysis were included. Enrichment significance was assessed using the Benjamini–Hochberg-adjusted p-values provided by Enrichr. For visualization, the ten most significantly enriched GO terms were displayed as dot plots generated with Seaborn, with dot size representing the number of overlapping proteins assigned to each GO term and color indicating the adjusted p-value.

### Structural modeling

Structural models of LRRK2–nanobody complexes were generated using the AlphaFold3 server (DeepMind/Isomorphic Labs). To increase prediction confidence, modeling was restricted to the smallest LRRK2 fragments that had previously been experimentally validated to bind the respective nanobodies by biochemical assays and cross-linking mass spectrometry (Singh *et al*., 2022). For the WD40-directed nanobodies Nb22 and Nb40, the kinase–WD40 fragment was used as input, whereas the full-length LRRK2 was used for Nb17. Predicted LRRK2-Nb22 and Nb40 complexes were superimposed onto the cryo-EM structure of full-length autoinhibited human LRRK2 (PDB: 7LHW) using PyMOL (Schrödinger, LLC) to assess compatibility with the experimentally determined conformation and to identify potential steric clashes. The selected AlphaFold3-derived Nb22 and Nb17 complex models were subjected to pose-preserving water refinement using the HADDOCK2.5 web server.

### Protein interaction network analysis

Protein interaction networks were generated from proteins identified as differentially abundant according to the statistical and effect-size criteria described above using an in-house Python workflow. For integrated analyses, proteins quantified in both corresponding comparisons were retained when they met the active fold-change and q-value criteria in at least one comparison (Nb22 versus Nb40 or Nb22 versus Nb17 for the nanobody dataset; MLi-2 versus DMSO or MLi-2 versus GZD-824 for the inhibitor dataset), while quantitative effects from both comparisons were retained for visualization. Protein associations were retrieved from a local IntAct database (PSI-MITAB 2.7, release 252, January 2026) and the STRING physical-interaction network (v12.0; accessed September 23, 2026) (Del Toro *et al*, 2022; Szklarczyk *et al*, 2025). IntAct interactions with MIscore ≥ 0.35 and STRING physical associations with confidence ≥ 0.40 were retained and weighted according to MIscore and STRING confidence, respectively. Where a protein pair was supported by both databases, the two sources were retained as independent evidence rather than combined into a single score. Network organization was determined exclusively from the retained confidence-weighted interaction topology and was independent of differential-abundance values or functional annotation. Connectivity-defined subnetworks were identified using a weighted Leiden-style community decomposition following the principles of the Leiden algorithm (Traag *et al*, 2019). Inter-community connectivity was further characterized using weighted edge betweenness and neighborhood Jaccard overlap; edge-betweenness approaches for identifying community structure have been described previously (Girvan & Newman, 2002). Node area was scaled according to the absolute log2 fold change, whereas node fill represented the direction and magnitude of differential abundance; community membership was indicated by categorical node-ring colors and hulls. LRRK2 was retained as the experimental bait using its measured quantitative values when available. For module-level visualization, topology-defined communities were coarse-grained using only retained protein–protein evidence: each unique protein pair contributed once, module area represented protein count, and inter-module edge width represented the summed weight of unique protein-pair connections; visualization-only scaffold edges were excluded. Following topology-based community assignment, communities were functionally annotated post hoc using Reactome release 97 (accessed September 23, 2026). Pathway over-representation was evaluated against proteins in the displayed network using a hypergeometric test with Benjamini–Hochberg correction within each community. For visualization, pathways with q ≤ 0.05 and an overlap of at least three proteins were retained, with up to two pathways displayed per community. Reactome annotations were used exclusively for biological interpretation and did not influence network construction, edge weighting, node coordinates, or community assignment.

### Downstream analysis of the two processed proteomic comparisons – trend analysis

The processed protein-level output tables from the Nb22-versus-Nb17 and Nb22-versus-Nb40 comparisons were used as the starting point for downstream integrative analysis. No additional normalization or missing-value imputation was performed. Because the pairwise comparisons were processed independently, missing values in the shared Nb22 condition could be imputed differently despite identical upstream settings. Shared Nb22 LFQ values were therefore compared between the processed output tables as a quality-control step. For proteins present in both datasets, protein-level Nb22 means from the two comparisons were averaged before construction of the three-state matrix.

For data-driven family discovery, the Nb22-versus-Nb17 comparison was used as the discovery contrast. Proteins were ranked using a signed significance score defined as sign(log2FC) × −log10(P), based on the Rubin-pooled differential-abundance statistics exported by the upstream analysis. The upper and lower 10% of the ranked distribution were defined as Nb22-high and Nb22-low tails, respectively. Candidate protein families were generated from shared leading gene-symbol prefixes and tested for over-representation in either tail using one-sided Fisher’s exact tests, with all ranked proteins as background. P values were adjusted using the Benjamini-Hochberg procedure. Candidate families were retained when at least three members occurred in the corresponding ranked tail, the enrichment odds ratio was ≥3, and the FDR-adjusted q value was <0.05. Exact redundant family calls identifying the same ranked-tail protein set were collapsed, and a nonredundant set of biologically interpretable families was fixed before ordered trend analysis. Representative anchor proteins were defined separately and were not preferentially weighted in module-level statistics.

In addition to the data-derived families, two exploratory groups were evaluated using the same trend-analysis framework. A cytosolic-proteostasis set comprised chaperone, redox, and peptidyl-prolyl isomerase proteins. A mitochondrial-surface set comprised cytosol-accessible mitochondrial proteins selected on the basis of subcellular localization together with positive +biotin-versus−biotin proximity support. These groups were not generated by the gene-prefix family-enrichment procedure and were therefore considered exploratory.

Independent +biotin-versus−biotin comparisons were used as an orthogonal assessment of proximity-labeling specificity. Positive biotin-dependent enrichment with FDR-adjusted q < 0.05 in at least two of the three nanobody conditions was used to identify specificity-supported proteins and to assess the robustness of module trajectories. YWHA/14-3-3 proteins were retained as a predefined exception in the specificity-supported sensitivity analysis because of the well-established role of 14-3-3 proteins in LRRK2 regulation.

For ordered trend analysis, proteins present in both processed pairwise comparison tables were retained. Mean LFQ abundance was calculated across the six measurements for Nb17 and Nb40. For Nb22, protein-level means from the two independently processed comparisons were averaged. Each protein was subsequently centered by subtracting its mean abundance across Nb17, Nb40, and Nb22. All quantified members of each fixed protein group were included.

Ordered changes across Nb17, Nb40, and Nb22 were evaluated using Page’s L trend test, treating proteins within each group as blocks and nanobody conditions as ordered states. For groups decreasing toward Nb22, the condition order was reversed for calculation of the one-sided Page statistic. P values were corrected across the tested groups using the Benjamini-Hochberg procedure. As an additional measure of group coherence, proteins displaying the strict monotonic ordering Nb17 < Nb40 <Nb22 or Nb17> Nb40 > Nb22 were counted according to the observed group direction. Enrichment of the specified strict ordering was assessed using a one-sided binomial test with an expected probability of 1/6, corresponding to one of the six possible strict orderings of three conditions.

For visualization, mean protein-centered log2 LFQ abundances were plotted across Nb17, Nb40, and Nb22. Shaded areas represent bootstrap 95% confidence intervals of the group mean obtained from 10,000 resamplings of proteins within each group. These intervals describe variation across proteins within a group and do not represent biological-replicate confidence intervals. All downstream analyses and graphical output were generated in Python.

## Generation of Images

Figures containing schematic illustrations and biological icons were created using BioRender (BioRender.com).

## Author contributions

B.R. and F.H. performed the experiments, analyzed the data, and contributed to manuscript preparation. F.H. K.B. A.R. F.R. and C.J.G. analyzed the data and wrote the manuscript. C.J.G. F.R. and W.V. conceived the study and developed the scientific concept. W.V. and C.G. provided the nanobody reagents and contributed to data analysis. All authors reviewed and approved the final manuscript.

## Data availability

The mass spectrometry proteomics data have been deposited to the ProteomeXchange Consortium via the PRIDE (Perez-Riverol *et al*, 2022) partner repository with the dataset identifiers PXD080701 and PXD080743. The datasets will become publicly available upon publication of the accepted manuscript.

## Acknowledgments

We are grateful to the staff of the Core Facility for Medical Proteomics at the Medical Faculty of the University of Tübingen for technical assistance. C.J.G. and W.V. received funds from The Michael J. Fox Foundation for Parkinson’s Research (MJFF-14527, MJFF-14527.01 and MJFF-023920). W.V. received funds from the VUB (SRP95 and OZR4292) and the Research Foundation Flanders (grant G031324N).

## Supplemental Figures

**Supplementary Figure 1:**
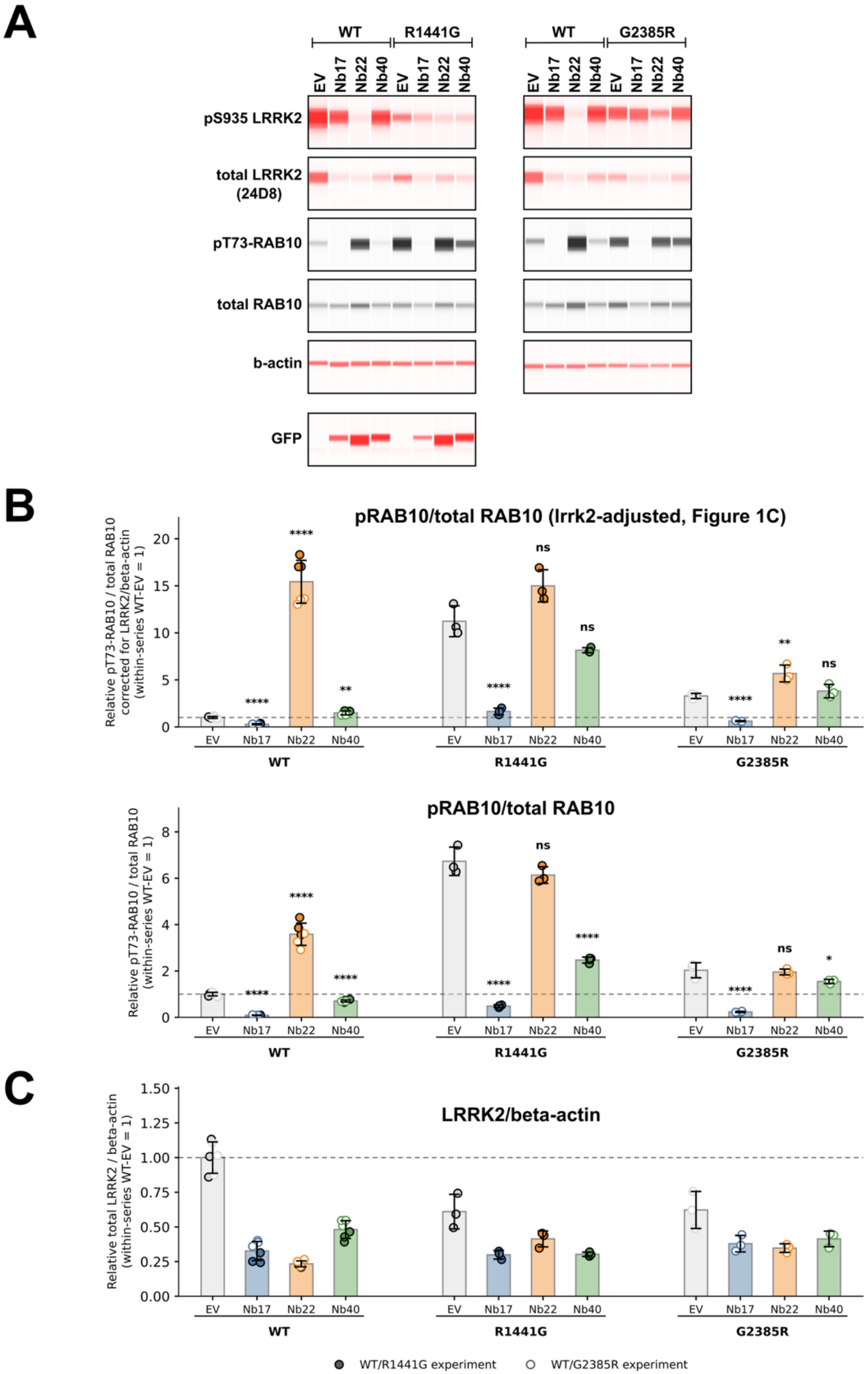
(A) Representative capillary Western blot images corresponding to the quantitative analyses shown in Figures 1B and 1C. pS935 and total-LRRK2 (24D8) were detected in the NIR channel, pT73-RAB10 and total-RAB were detected via chemiluminescence. GFP and b-actin controls were detected in the NIR channel. Left panel: WT/R1441G comparison; right panel: WT/G2385R comparison. (B) Side-by-side comparison of LRRK2-adjusted pRAB10 levels shown in Figure 1C (upper panel) to pRab levels normalized to total-RAB10 only (lower panel). (C) relative LRRK2 expression levels (normalized to β-actin) Values were normalized to the WT-EV (empty vector) control within each experimental series (WT = 1). Individual biological replicates are shown as circles, with horizontal lines indicating the mean and error bars representing SD. Statistical significance in both panel (B) was determined by genotype-wise Dunnett-adjusted multiple comparisons versus the EV control following within-series normalization. ns, P ≥ 0.05; P < 0.05 (*); P < 0.01 (**); P < 0.001 (***); P < 0.0001 (****). Each mutant condition comprises three biological replicates. WT data from the WT/R1441G and WT/G2385R experimental series were combined after normalization to the corresponding within-series WT-EV control (WT = 1), yielding six observations for each WT condition.

**Supplementary Figure 2:**
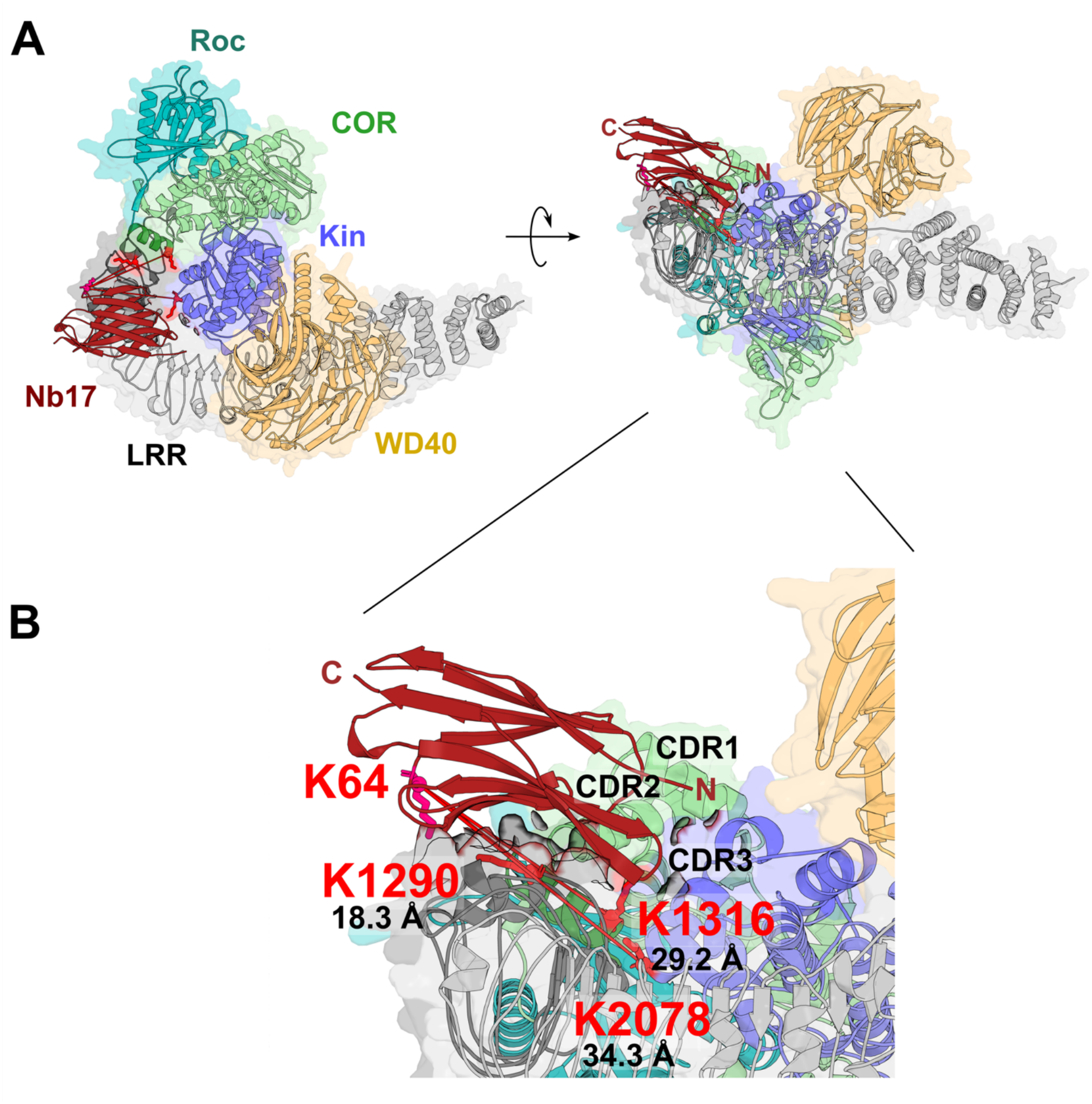
AF3-derived model of the Nb17–LRRK2 binding interface. (A) Cartoon and semitransparent surface representations of the AF3-derived Nb17–LRRK2 model, trimmed to the LRRK2 residues resolved in the experimental structure 7LHW. The N- and C-terminal portions of the LRR region are colored light and dark gray, respectively; the α0 helix is forest green, ROC teal, COR pale green, the kinase domain slate blue, and WD40 light orange. Nb17 is shown in dark red. The DSSO crosslinks reported by Singh et al. (2022) are indicated, with the crosslinked residues highlighted in red. The model positions Nb17 predominantly against the C-terminal LRR surface, with an additional contribution from the kinase C-lobe. (B) Enlarged view of the predicted interface. The Cα–Cα distances from Nb17 K64 to LRRK2 K1290, K1316 and K2078 are 18.3, 29.2 and 34.3 Å, respectively. All three are compatible with the relaxed 35 Å distance threshold commonly applied to DSSO crosslinks when protein dynamics are considered. Residues from all three Nb17 CDR loops contribute to the modeled interface. This crosslink-compatible pose provides a structural hypothesis in which Nb17 bridges the LRR and kinase regions and thereby could stabilize a closed, inhibited LRRK2 conformation.

**Supplementary Figure 3:**
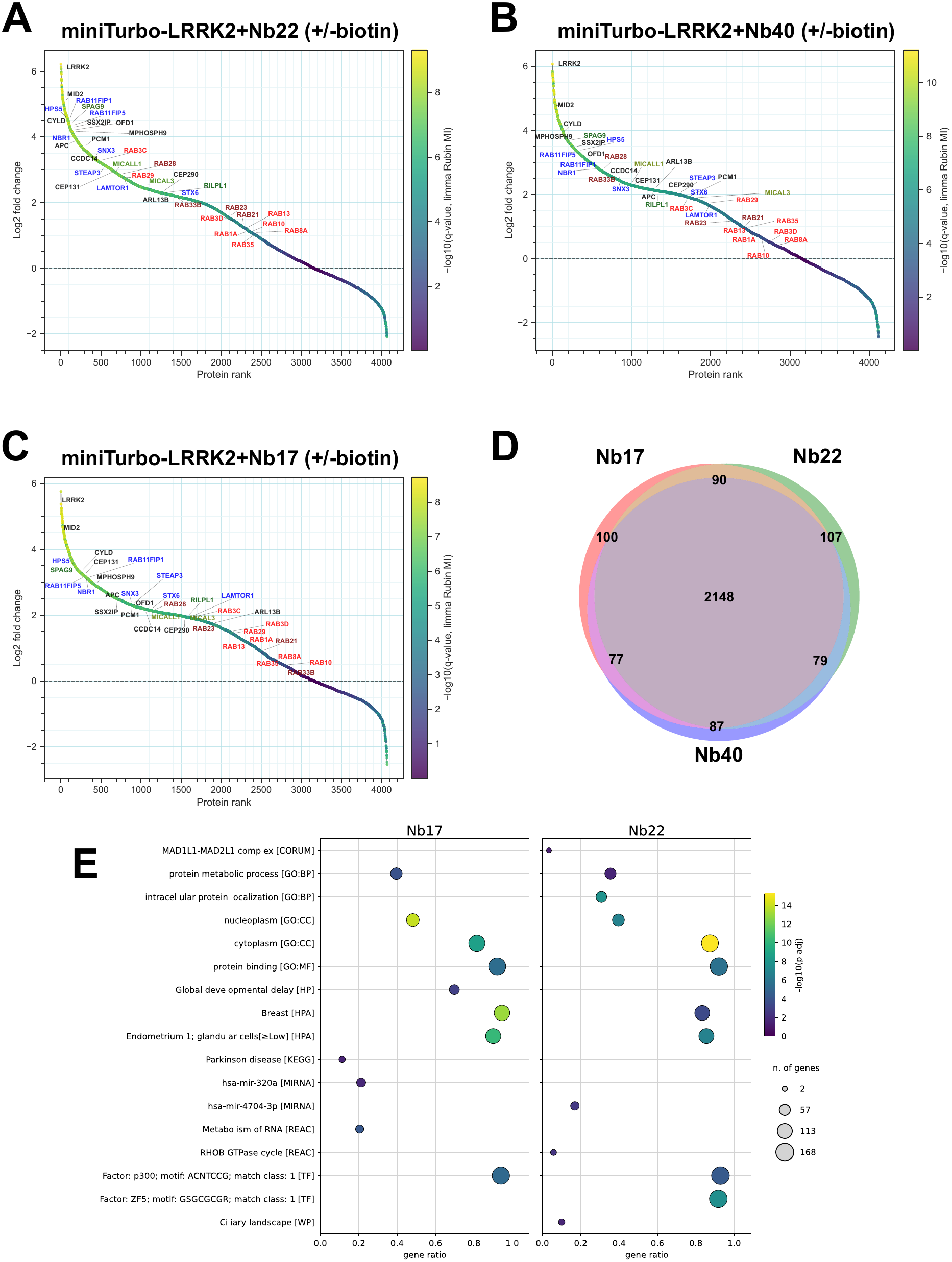
Biotin-dependent enrichment of the LRRK2 proximity proteome. Ranked protein plots showing the log₂ fold change between biotin-labeled (+biotin) and unlabeled (−biotin) samples following miniTurbo-mediated proximity labeling of LRRK2. Proteins are ranked according to their biotin-dependent enrichment, with the y-axis indicating the log₂ fold change (+biotin/−biotin). Point color represents statistical significance (−log₁₀ adjusted q-value) derived from the Rubin-pooled LIMMA analysis. Selected LRRK2-associated proteins are annotated to illustrate robust biotin-dependent enrichment. Rab GTPases are color-coded in red and brown, with established LRRK2 Rab substrates shown in red. Rab effector proteins are shown in green, with established downstream effectors of LRRK2 highlighted in dark green. (A) miniTurbo-LRRK2 co-expressed with the activating nanobody Nb22. (B) miniTurbo-LRRK2 co-expressed with the neutral nanobody Nb40. (C) miniTurbo-LRRK2 co-expressed with the inhibitory nanobody Nb17. (D) Venn diagram showing the overlap of the three conditions (Nb17, Nb22 and Nb40). (E) Enrichment analysis showing top significantly enriched GO-terms in the significant proteins exclusively found in the Nb17 and the Nb22 condition.

**Supplementary Figure 4:**
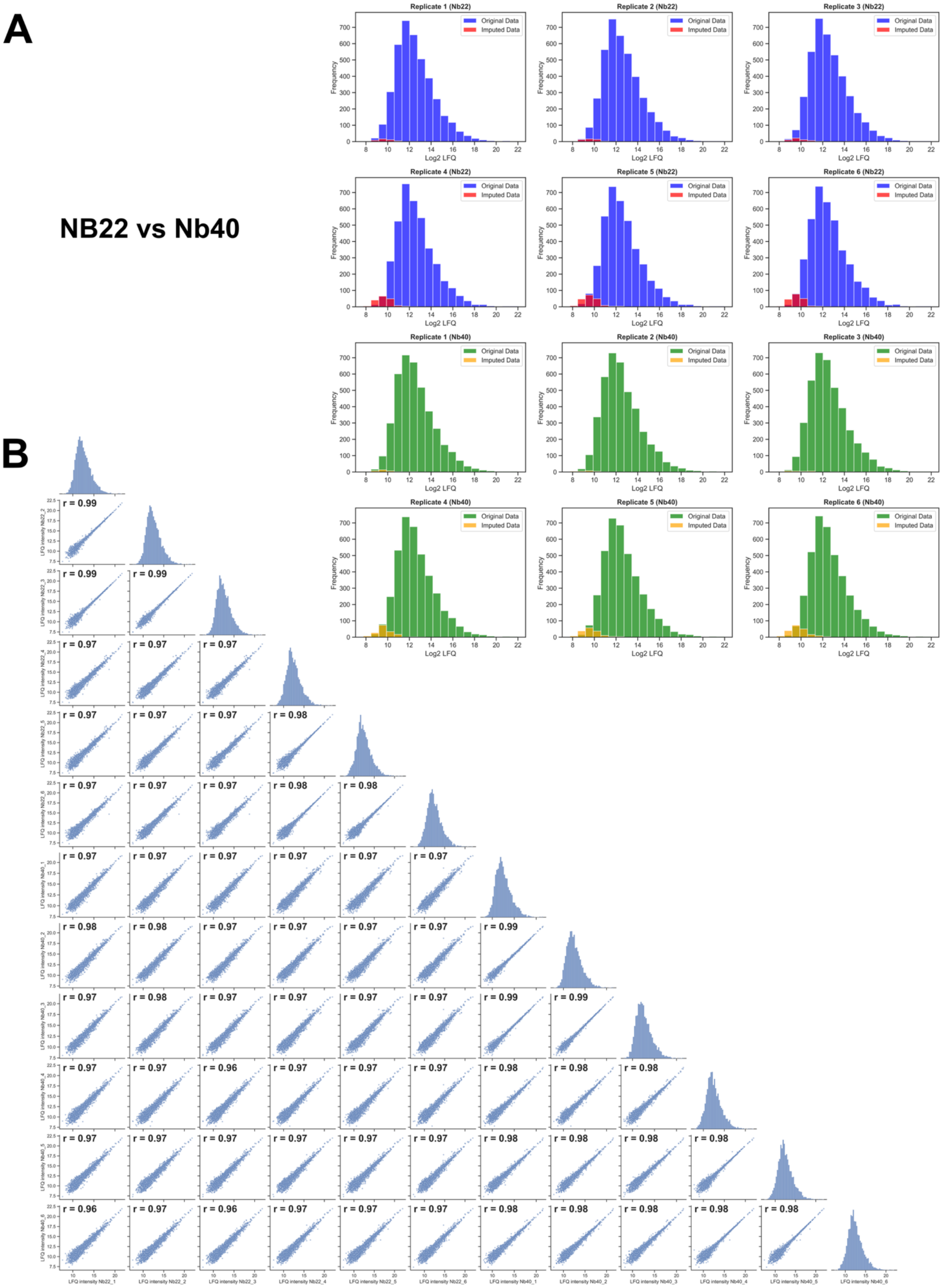
Comparison Nb22 vs Nb40. (A) Histograms for each condition. Effect of MNAR imputation is indicated. (B) Scatter plots comparing all conditions for quality assessment.

**Supplementary Figure 5:**
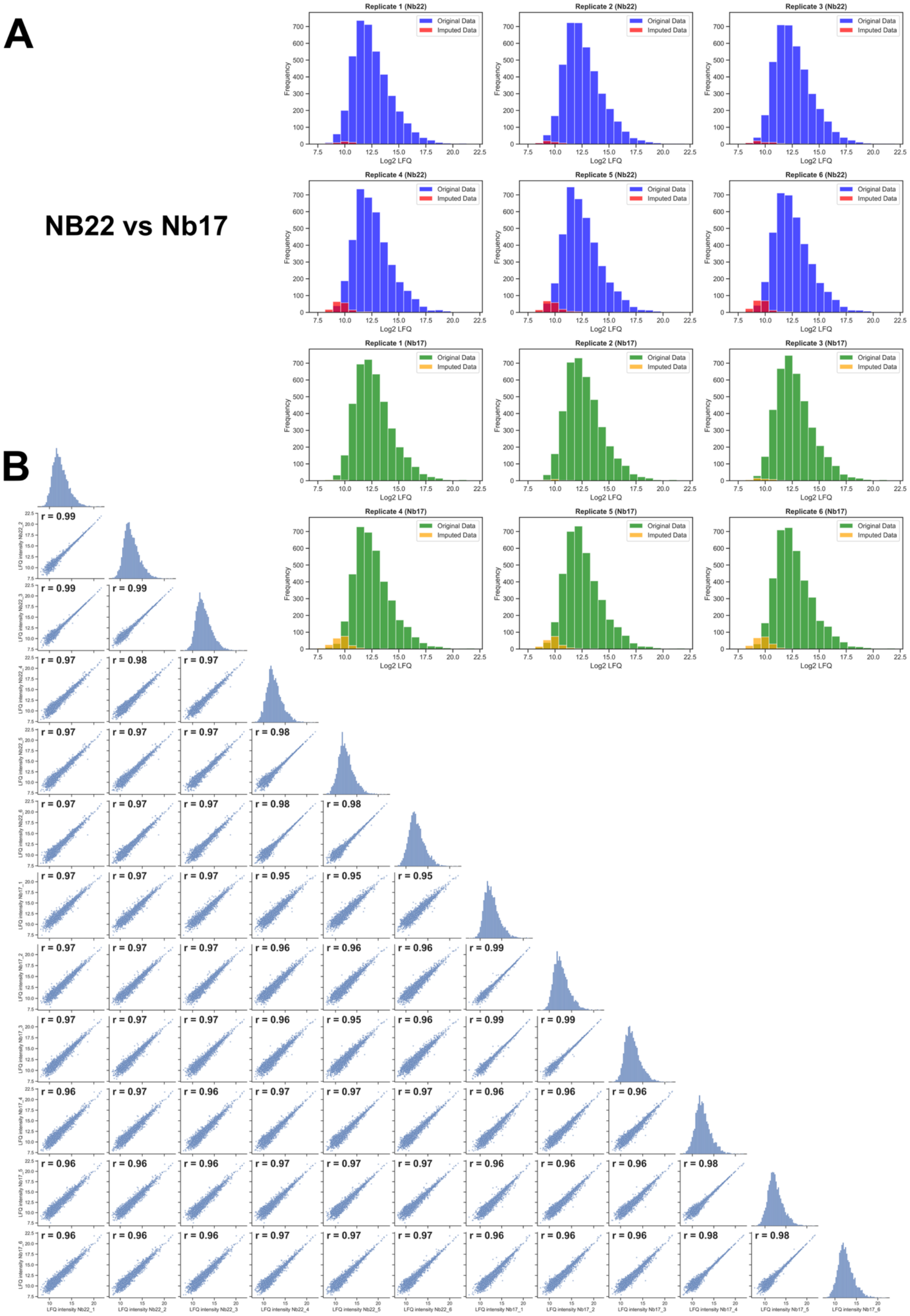
Comparison Nb22 vs Nb17. (A) Histograms for each condition. Effect of MNAR imputation is indicated. (B) Scatter plots comparing all conditions for quality assessment.

**Supplementary Figure 6:**
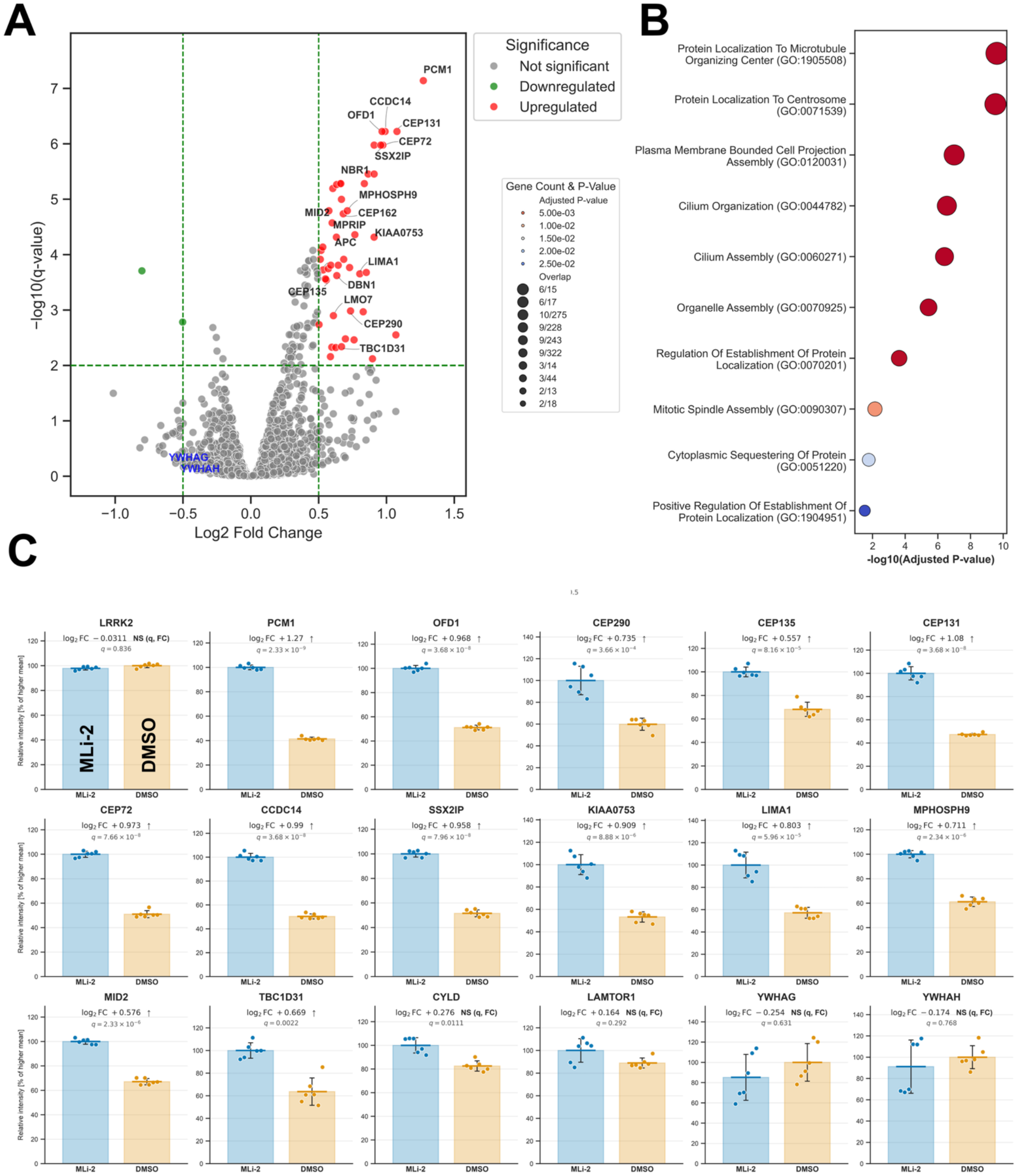
Rewiring of the LRRK2 proximity proteome by the type I inhibitor MLi-2 compared to a DMSO control replicating our previous data by DIA-PASEF (Eckert *et al*., 2026) at increased sensitivity. (A) Volcano plot showing differential protein abundance analyzed using a limma-like moderated t-test with multiple imputation and Rubin pooling; P values were adjusted using the Benjamini–Hochberg method (N=6 [2 biological and 3 technical replicates]; Rubin MI; FDR=0.01, N proteins=3768). (B) GO-term analysis of the proximity proteome enriched with MLi-2. (C) Pairwise comparison of selected proteins (mean normalized abundance N=6 [2 biological and 3 technical replicates]), error bar= SD, enriched protein set to 100%, LIMMA q-value is shown (ns= non-significant, up and down arrows indicate the direction of significant differences).

**Supplementary Figure 7:**
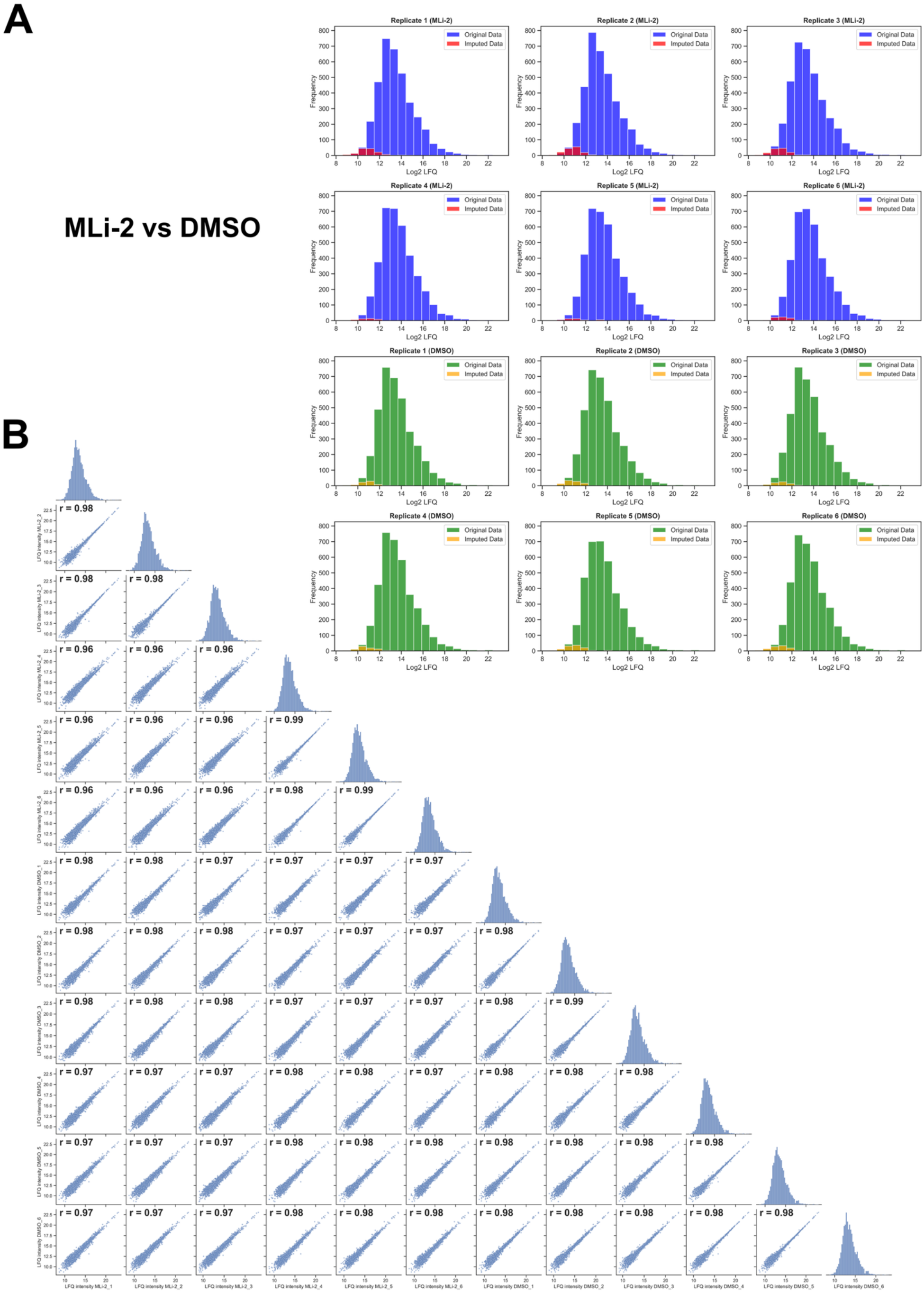
Comparison MLi-2 vs DMSO. (A) Histograms for each condition. Effect of MNAR imputation is indicated. (B) Scatter plots comparing all conditions for quality assessment.

**Supplementary Figure 8:**
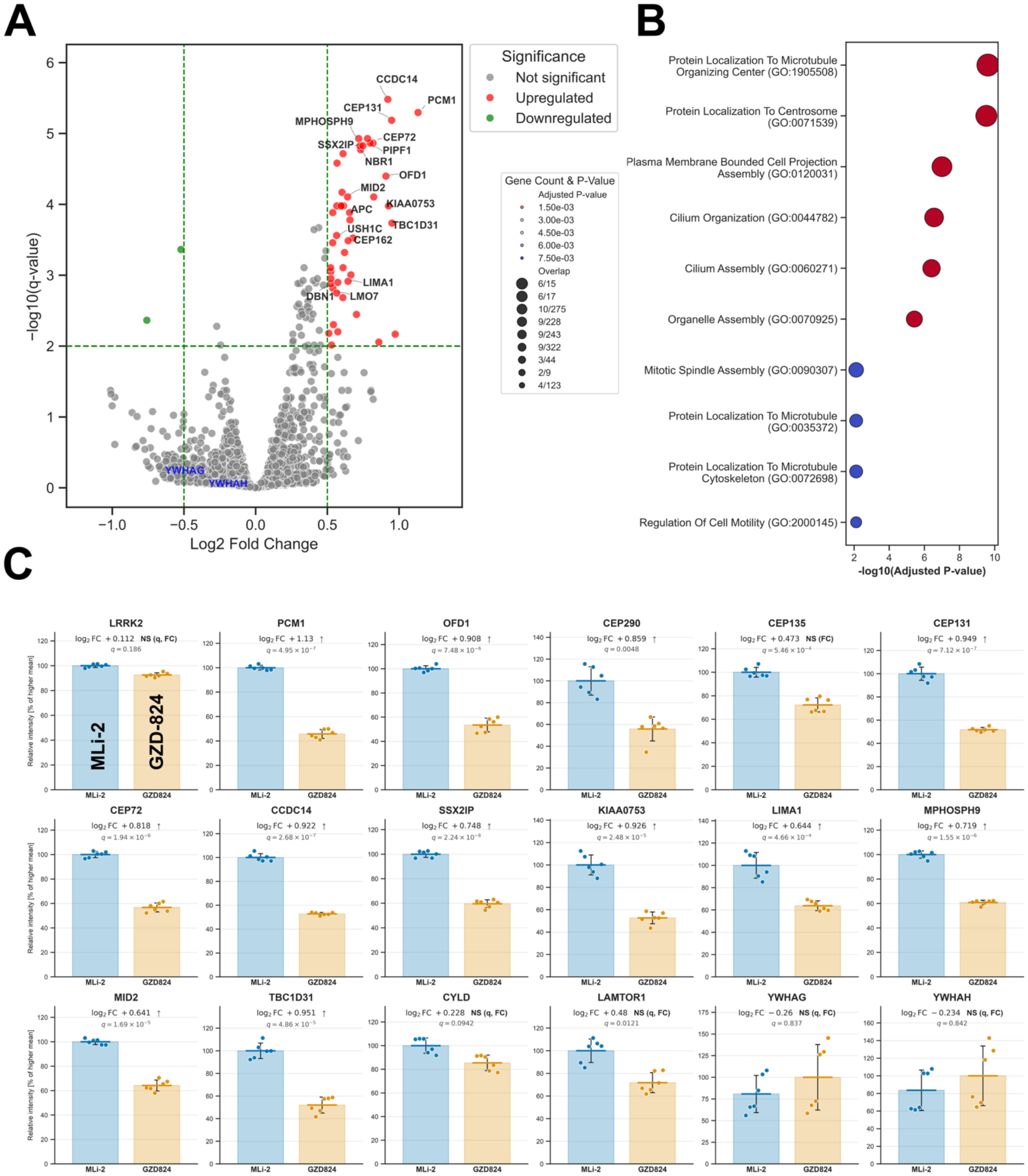
Rewiring of the LRRK2 proximity proteome by the type I inhibitor MLi-2 compared to the type II inhibitor GZD-824 replicating our previous data by DIA-PASEF (Eckert *et al*., 2026) at increased sensitivity. (A) Volcano plot showing differential protein abundance analyzed using a limma-like moderated t-test with multiple imputation and Rubin pooling; P values were adjusted using the Benjamini–Hochberg method (N=6 [2 biological and 3 technical replicates]; Rubin MI; FDR=0.01; N proteins=3834). (B) GO-term analysis of the proximity proteome enriched with MLi-2. (C) Pairwise comparison of selected proteins (mean normalized abundance N=6 [2 biological and 3 technical replicates]), error bar= SD, enriched protein set to 100%, LIMMA q-value is shown (ns= non-significant, up and down arrows indicate the direction of significant differences).

**Supplementary Figure 9:**
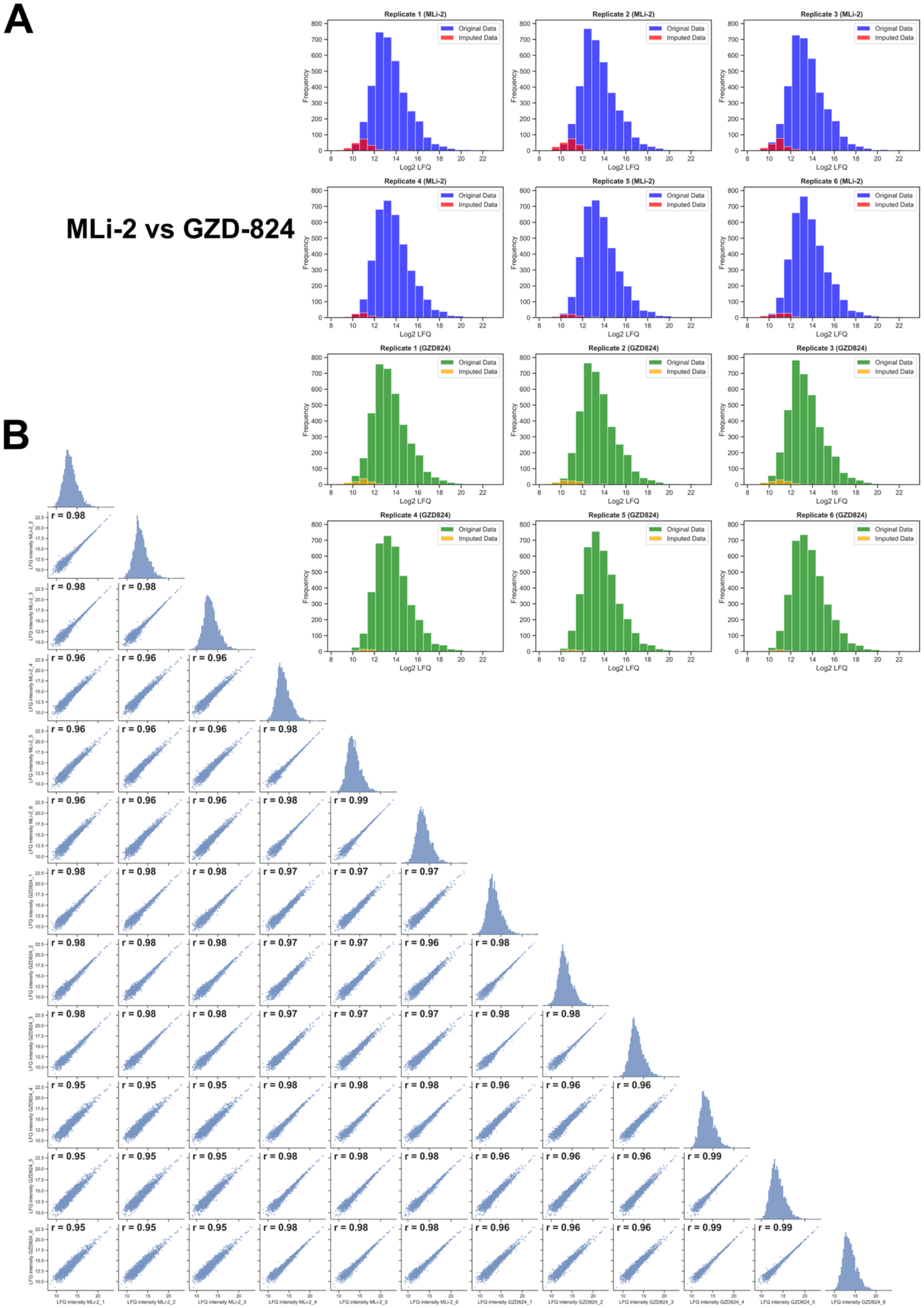
Comparison MLi-2 vs GZD-824. (A) Histograms for each condition. Effect of MNAR imputation is indicated. (B) Scatter plots comparing all conditions for quality assessment.

**Supplementary Figure 10:**
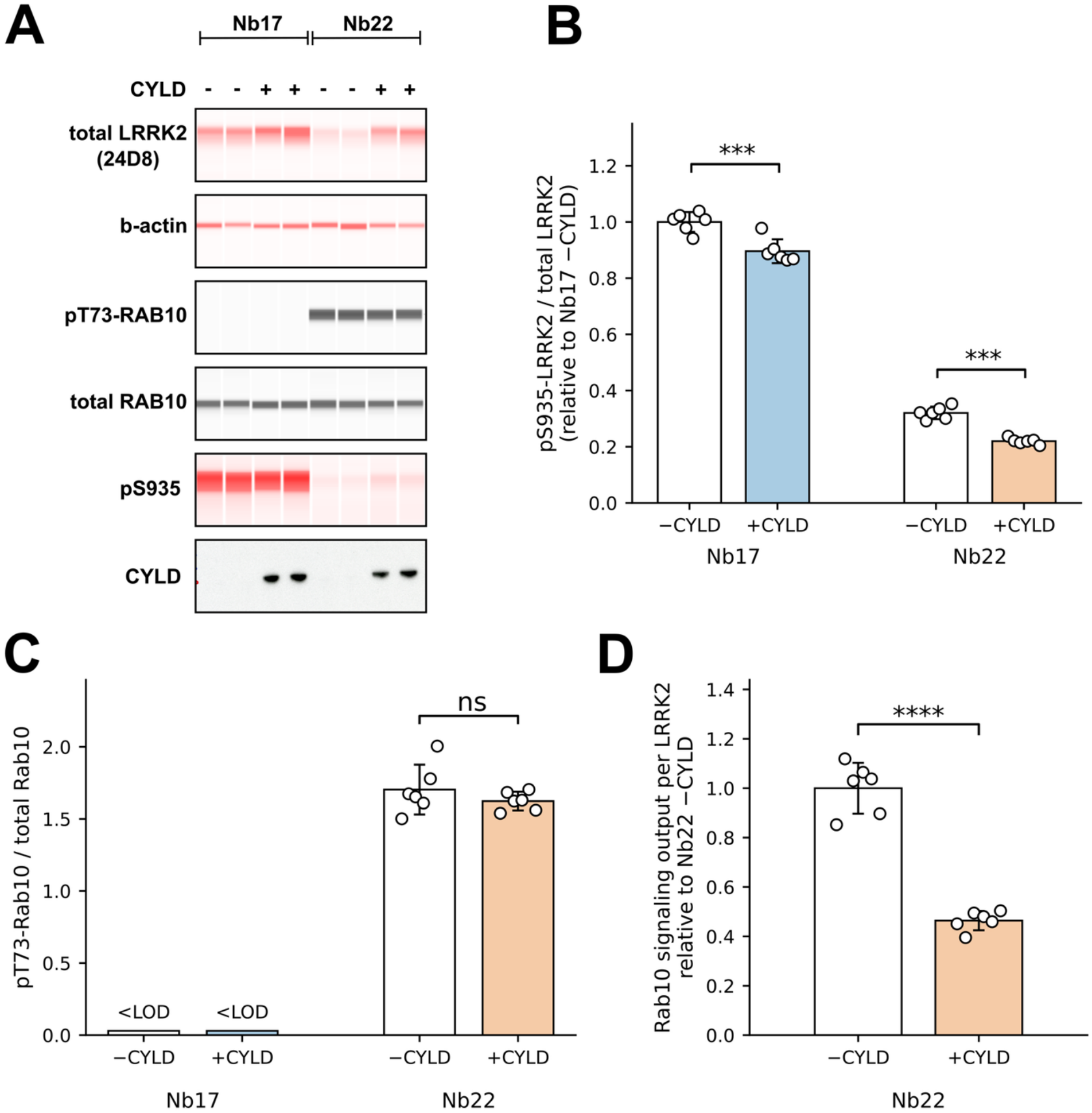
Effect of CYLD co-expression on LRRK2 abundance and Rab10 phosphorylation. (A) Representative capillary Western analyses of HEK293T cells expressing WT LRRK2 together with Nb17 or Nb22 in the absence (−CYLD) or presence (+CYLD) of co-expressed CYLD. Total LRRK2 (24D8), β-actin, pT73-RAB10, total RAB10, pS935-LRRK2, and CYLD were detected. Total LRRK2 and β-actin, and pT73-RAB10 and total RAB10, were analyzed on separate Jess arrays. CYLD (anti-CYLD) was detected by classical Western blot. (B) Quantification of pS935-LRRK2 phosphorylation normalized to total LRRK2 and expressed relative to the mean of the Nb17/−CYLD condition. CYLD co-expression resulted in a modest but significantly reduced relative pS935 phosphorylation in both the Nb17 and Nb22 conditions. (C) Quantification of pT73-Rab10 relative to total Rab10. In Nb17-expressing cells, pT73-Rab10 was below the limit of detection and was therefore not subjected to quantitative statistical analysis. CYLD co-expression did not significantly alter pT73-Rab10/total Rab10 in the Nb22 condition. (D) Rab10 phosphorylation in the Nb22 condition additionally normalized to the corresponding β-actin-normalized LRRK2 abundance and expressed relative to the mean of the −CYLD condition. Bars represent mean ± SD and individual data points represent biological replicates (N = 6). Statistical significance was assessed separately for the effect of CYLD within the Nb17- and Nb22-bound LRRK2 conditions using two-sided unpaired Student’s t-tests (ns, P ≥ 0.05; *** P < 0.001; **** P < 0.0001).

**Supplementary Figure 11:**
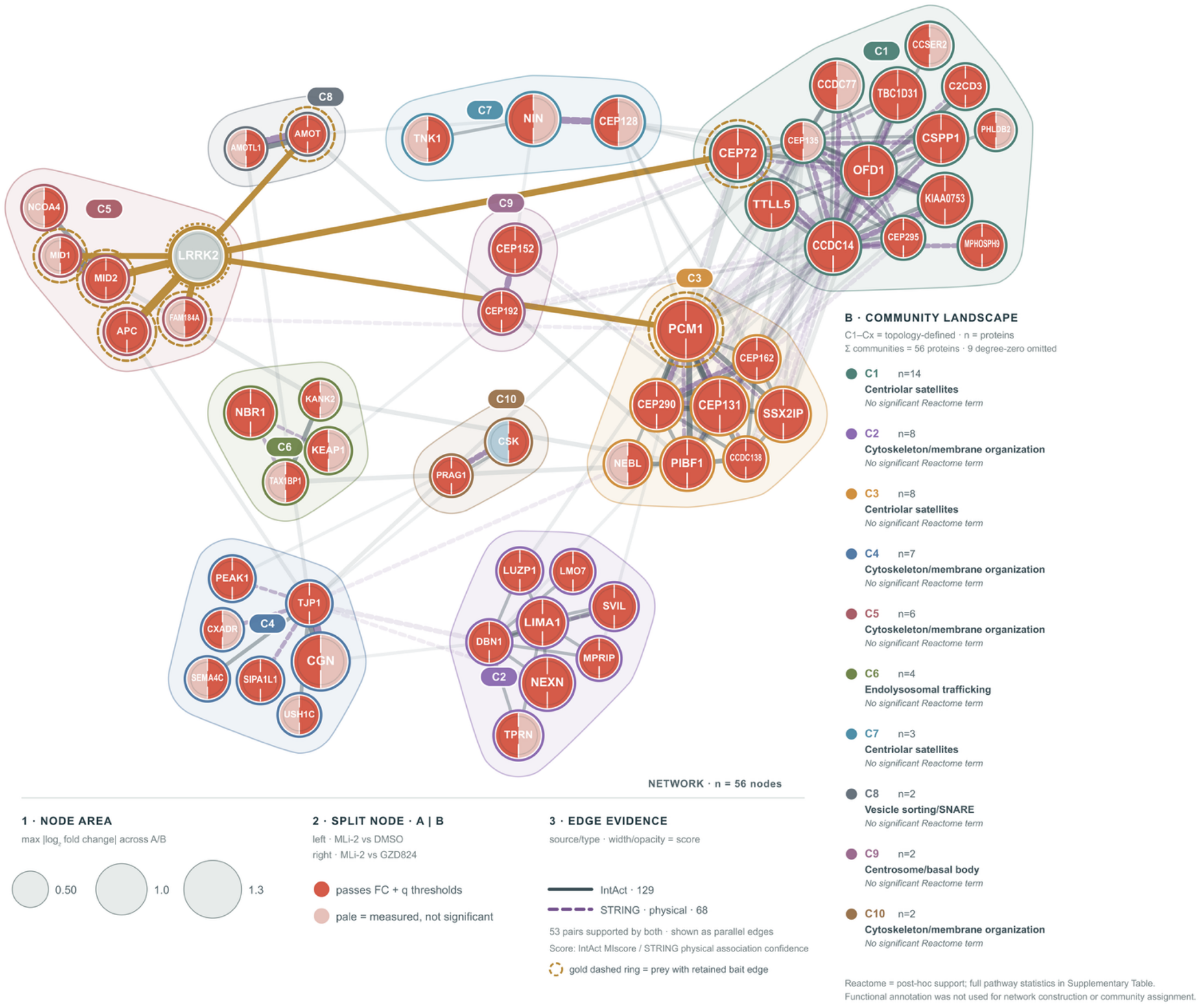
Comparison of LRRK2 proximity-proteome remodeling induced by MLi-2. Direct comparison of protein abundance changes in MLi-2 versus DMSO and MLi-2 versus GZD-824. Proteins are plotted according to their log₂ fold changes in the two comparisons, illustrating the concordance of the MLi-2-associated proximity-proteome signature across the two reference conditions. The network comprises 56 connected nodes partitioned into 10 communities (C1–C10); 9 proteins without network connections are not shown.

